# NFkB-signaling promotes glial reactivity and suppresses Müller glia-mediated neuron regeneration in the mammalian retina

**DOI:** 10.1101/2021.10.05.463152

**Authors:** Isabella Palazzo, Levi J. Todd, Thanh V. Hoang, Thomas A. Reh, Seth Blackshaw, Andy J. Fischer

## Abstract

Müller glia (MG) in mammalian retinas are incapable of regenerating neurons after damage, whereas the MG in lower vertebrates regenerate functional neurons. Identification networks that regulate MG-mediated regeneration is key to harnessing the regenerative potential of MG. Here we study how NFkB-signaling influences glial responses to damage and reprogramming of MG into neurons in the rodent retina. We find activation of NFkB and dynamic expression of NFkB-associated genes in MG after damage, however NFkB activity is inhibited by microglia ablation. Knockout of NFkB in MG suppressed the accumulation of immune cells after damage. Inhibition of NFkB following NMDA-damage significantly enhanced the reprogramming of Ascl1-overexpressing MG into neuron-like cells. scRNA-seq of retinal glia following inhibition of NFkB reveals coordination with signaling via TGFβ2 and suppression of NFI and Id transcription factors. Inhibition of Smad3 or Id transcription factors increased numbers of neuron-like cells produced by Ascl1-overexpressing MG. We conclude that NFkB is a key signaling hub that is activated in MG after damage, mediates the accumulation of immune cells, and suppresses the neurogenic potential of MG.

## Introduction

The phenomenon of retinal regeneration has been studied for many decades. Within the last 2 decades Müller glia (MG) have been identified as the primary cellular source of retinal regeneration^1–4^. In normal healthy retinas MG are the primary support cells providing structural, metabolic, and synaptic support^5^. However, after damage MG can be stimulated to become activated, de-differentiate, proliferate as progenitor-like cells and produce progeny that differentiate into functional neurons that restore vision^1,4,6–8^. However, the neurogenic capacity of MG varies widely across species^9,10^. In lower vertebrate species, such as zebrafish, MG have an extraordinary capacity to become proliferating progenitors that regenerate functional neurons^1,3,11^. In birds, MG are capable of forming numerous proliferating progenitor-like cells with limited neurogenic potential^1^. By contrast, MG in the mammalian retina lack a significant regenerative response and, instead, rapidly activate a gliotic and quiescence-restoring programs^12^.

Comparative epigenomic and transcriptomic analysis of MG revealed that following damage, zebrafish and chick MG transition to a reactive state prior to reprogramming into progenitor cells whereas mammalian MG enter a reactive state prior to restoring quiescence^13^. NFkB has been implicated as a master regulator of inflammation^14^, and may be a key difference between species wherein components of this pathway are prevalently expressed in mammalian MG but to a lesser degree in MG chick^15^ and fish retinas^13^. Thus, we hypothesize that NFkB signaling may be part of the regulatory networks that drive reactive MG to restore quiescence and suppress gene modules that promote the de-differentiation and the formation of neurogenic progenitors. We have recently reported that NFkB signaling in the chick retina suppresses the formation of MGPCs and this process depends on signals from reactive microglia^15^. NFkB signaling is activated in the retina following different types of neuronal damage^16–18^ or chronic degeneration ^19,20^. NFkB promotes inflammation and exacerbates cell death in the mammalian retina^16,17^. However, nothing is known about how this pathway influences the ability of MG to reprogram into progenitor cells and generate neurons in the mammalian retina.

Although mammalian MG do not spontaneously undergo reprogramming into neurogenic progenitor-like cells in response to retinal damage, overexpression of Ascl1 in combination with damage stimulates the reprogramming of MG into functional neurons^21^. It was recently reported that Jak/Stat signaling^22^ and signals produced by reactive microglia suppress neuronal regeneration from Ascl1-overexpressing MG ^23^.

The purpose of this study was to investigate how NFkB influences glial responses to retinal damage and Ascl1-mediated reprogramming of MG. We find that NFkB is rapidly activated in MG and that is dependent on signals from microglia. Further, we find that NFkB signaling interferes with Ascl1-mediated neuronal regeneration by promoting expression of pro-glial transcription factors that drive MG to restore quiescence.

## Results

### scRNA-seq analysis of NFkB-related genes in damaged mouse retina

We first assessed the patterns of expression of NFkB-related factors across different cell types in normal and damaged mouse retinas. We analyzed single cell RNA sequencing (scRNA-seq) libraries from control and NMDA-damaged WT retinas^13^. Uniform Manifold Approximation and Projection (UMAP) plots revealed discrete clusters of different retinal cell types **(Fig. 1a)**. Cell clusters were labeled based on gene expression patterns described in the Methods. Neuronal cells from control and damaged retinas were clustered together regardless of time after NMDA-treatment **(Fig. 1a)**. By contrast, resting MG, including cells from 48 and 72 hrs after NMDA, and activated MG from 3, 6, 12, and 24 hrs after NMDA were spatially separated in UMAP plots **(Fig.1a,b)**. Pseudotime analysis generated a trajectory of cells with resting MG (control MG and some MG from 48 to 72 hr after treatment) to the left, MG from 3 to 6 hr after treatment to the far right, and MG from 12 to 24 hr bridging the middle **(Supplemental Fig. 1a–d)**.

**Figure 1:**
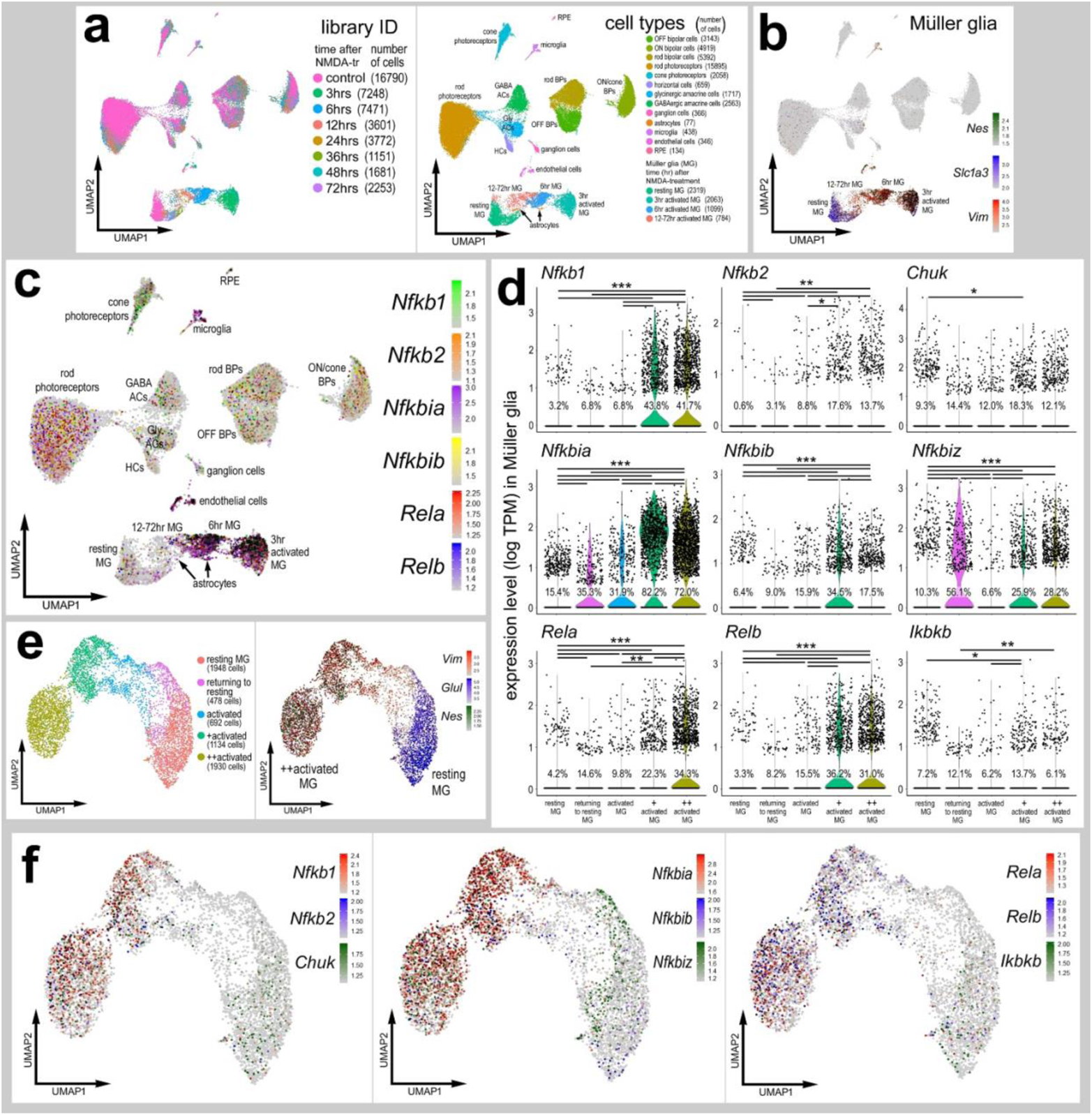
scRNA-seq analysis of NFkB-related genes in damaged mouse retina. UMAPs of aggregated scRNA-seq libraries prepared from whole wild type control retinas and retinas 3hr, 6hr, 12hr, 24hr, 36hr, 48hr, and 72hr after NMDA damage; numbers of cells per library or per cluster in parentheses **(a)**. Clusters of different types of retinal cells were identified based on collective expression of different cell-distinguishing markers as described in the Materials and Methods. Resting MG and reactive MG identified by expression of *Slc1a3* or *Nes and Vim*, respectively **(b)**. UMAP plots illustrate expression of *Nfkb1, Nfkb2, Nfkbia, Nfkbib, Nfkbiz, Rela, Relb, Ikbkb*, and *Chuk* **(c)**. Violin/scatter plots illustrate expression levels of these factors within MG **(d)**. Significance of difference (**P<0.01, ***P<0.001) was determined by using a Wilcoxon rank sum with Bonferroni correction. UMAP plots of isolated and re-embedded MG with activation states identified by expression of *Glul, Vim*, and *Nes* **(e)**. UMAP plots of MG illustrate expression of *Nfkb1, Nfkb2, Chuk, Nfkbia, Nfkbib, Nfkbiz, Rela, Relb, Ikbkb* **(f)**.

We examined changes in expression levels of NFkB-related genes including transcription factors *Nfkb1* (p105/50), *Nfkb2* (p100/52), *Rela* (p65), and *Relb*, inhibitor of kappa-B (IkB) components *Nfkbia* (IkB-a), *Nfkbib* (IkB-b), *Nfkbiz* (IkB-z), and inhibitor of kappa-B kinase (IKK) component *Chuk* (IKK-alpha). Although some NFkB-related genes were detected in different types of retinal neurons, most NFkB-related genes were highly expressed in endothelial cells, microglia, astrocytes and MG **(Fig. 1c)**. In microglia, for example, *Nfkb1, Nfkb2* and *Nfkbia/b/z* were significantly upregulated shortly after NMDA-treatment **(Supplemental Fig. 2)**. We next isolated the MG and identified significant increases in expression levels of 9 different NFkB-related genes in activated MG at 3 and 6hrs after NMDA treatment compared to levels seen in resting and undamaged MG **(Fig. 1d-f)**. These data suggest that genes involved in NFkB signaling are rapidly and highly upregulated in MG after damage. Dynamic changes of mRNA levels are strongly correlated with changes in protein levels and function^24^.

**Figure 2:**
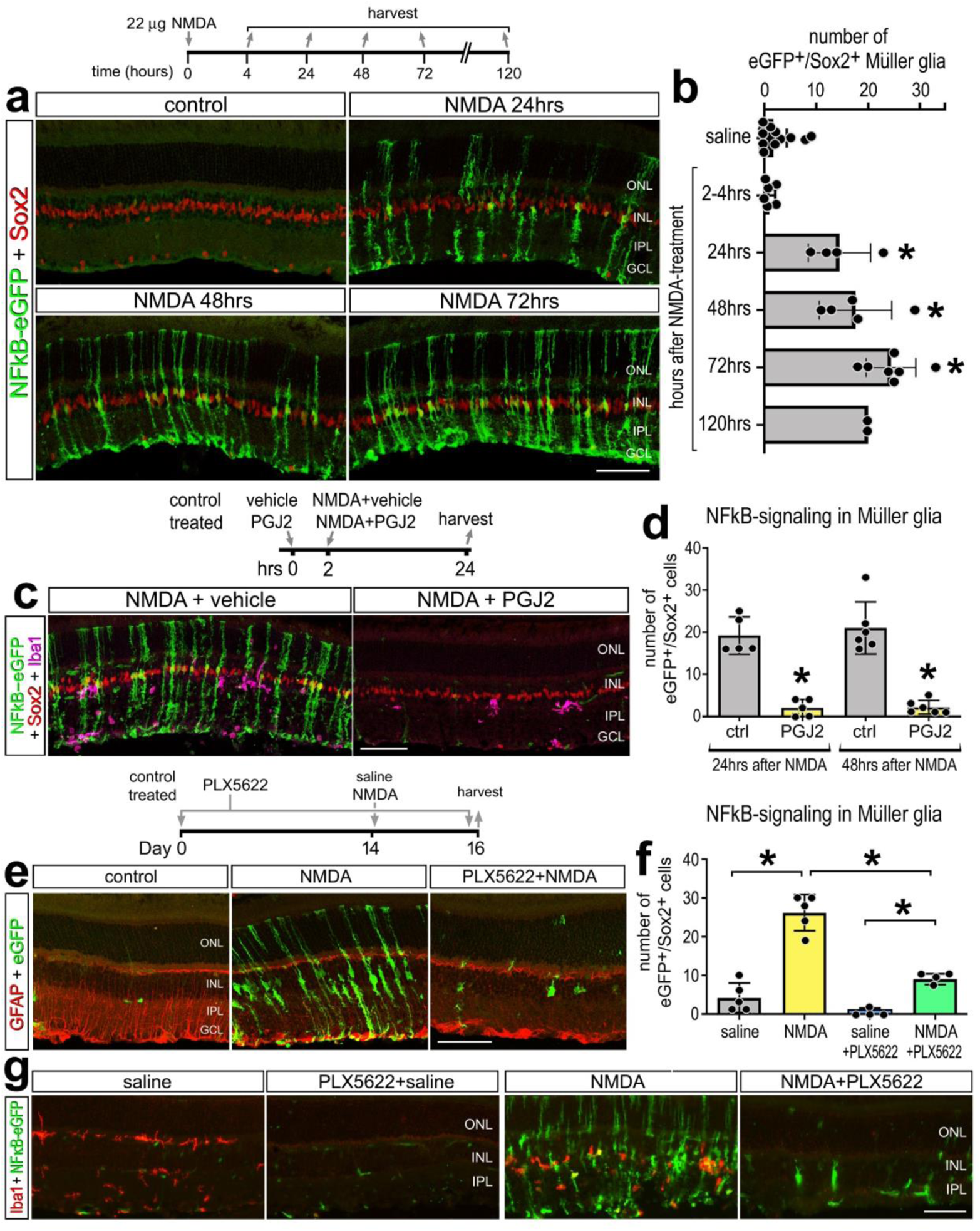
NFkB activation in response to retinal damage is dependent on microglia. Representative images of undamaged retinas and retinas 24hr, 48hr, or 72hr after NMDA damage from NFkB-eGFP reporter mice. Retinal sections were immunolabeled for GFP (green) and Sox2 (red) **(a)**. The histogram in **(b)** illustrated the mean (±SD and individual data points) numbers of Sox2^+^/GFP^+^ cells per field of view. Eyes were injected with vehicle control (left eye) or PGJ2 (NFkB inhibitor; right eye) 2h prior to NMDA treatment. Retinal sections were labeled for GFP (green), Sox2 (red) and Iba1 (magenta) **(c)**. The histogram in **(d)** illustrates mean (±SD and individual data points) numbers of Sox2^+^/GFP^+^ cells per field of view. Mice were fed a control diet or PLX5622-diet for 2 weeks, eyes were injected with saline or NMDA, and retinas harvest 2 days later (**e-g**). Retinal sections were immuno-labeled for GFP (green, **e,g**), GFAP (red, **e**), or Iba1 (red, **g**). Significance of difference (*p<0.05, **p<0.001) was determined by using one way ANOVA and Tukey’s post-hoc test. The calibration bars in panels **a, c, e** and **g** represent 50 µm. Abbreviations: ONL – outer nuclear layer, INL – inner nuclear layer, IPL – inner plexiform layer, GCL – ganglion cell layer.

### NFkB activation in response to retinal damage is dependent on microglia

To validate findings from scRNA-seq analyses, we identified patterns of NFkB activation *in situ* in the retina. We injected NMDA into the vitreous chamber of transgenic mice wherein eGFP is under the transcriptional control of NFkB cis regulatory elements (NFkB-eGFP)^25^. We observed significant increases in numbers of GFP^+^/Sox2^+^ MG at 24, 48, and 72hrs after damage **(Fig. 2a-b)**. Numbers of reporter-positive MG remained elevated at 120hrs after NMDA-treatment, but this likely represented the perdurance of eGFP. At 2 weeks after treatment there were few reporter-positive MG (not shown). Although the NFkB reporter was prevalently expressed by a majority of MG, activation was not detected in astrocytes or microglia, but was observed in a few endothelial cells (not shown), despite scRNA-seq evidence for expression of some NFkB-related genes in these cell types **(Fig. 1c-d; Supplemental Fig. 2)**. The absence of NFkB-reporter in microglia may be due to the specific NFkB cis-regulatory elements used in this transgenic line or may represent limited NFkB signaling without widespread expression of all components of this pathway.

We next injected a small molecule inhibitor, 15-deoxy-delta-12,14-prostaglandin J2 (PGJ2), into NMDA damaged eyes to test the efficacy of this NFkB inhibitor. PGJ2 has been previously shown to effectively inhibit NFkB signaling in different cell lines and chick retina ^15,26^. We observed a significant decrease in numbers of GFP^+^/Sox2^+^ MG in damaged retinas treated with PGJ2 compared to controls **(Fig. 2c-d)**. NFkB can be activated by various proinflammatory cytokines^14,27^, including IL1α, IL1β and TNF which are rapidly upregulated by retinal microglia after damage^28^. Thus, we examined how elimination of microglia influenced damage-induced activation of the NFkB-reporter in the retina. Microglia were ablated using PLX5622, a Csf1r antagonist that ablates CNS microglia within 2 weeks of treatment ^29,30^. Following 2 weeks of PLX5622 administration, we injected NMDA and observed a large significant decrease in numbers of GFP^+^/Sox2^+^ MG compared to numbers seen in control damaged retinas with microglia present **(Fig. 2e-f)**. The efficacy of the PLX5622 induced ablation of microglia was confirmed by immunolabeling labeling for Iba1 **(Fig. 2g)**. Taken together, these data indicate that NFkB signaling is robustly activated in MG after damage and this activation can be blocked by small molecule inhibitors or ablation of microglia, implying that reactive microglia secrete factors that selectively activate NFkB signaling in MG.

### Conditional knockout of NFkB signaling in MG impairs immune cell responses in damaged retinas

We next examined how genetic deleteion of NFkB signaling in MG influences cell survival and the responses of immune cells. Upon NFkB activation, the IKK complex is activated and phosphorylates IkBa/IkBb, thereby releasing NFkB transcription factors and allowing them to translocate into the nucleus to regulate transcription of target genes^31^. Conditional knockout of *Ikkb* blocks signaling through the canonical NFkB pathway due to the loss of IKK-mediated phosphorylation and degradation of IkBa/IkBb, thereby leading to maintained sequestration of NFkB transcription factors in the cytoplasm^31,32^. To permit tamoxifen-inducible deletion of *Ikkb* in MG, mice with floxed alleles of *Ikkb* (*Ikkb^fl/fl^*)^32^ gene were crossed with *Rlbp1-CreERT* mice (*Rlbp1-CreERT:Ikkb^fl/fl^*). We administered 4 consecutive daily doses of tamoxifen prior to NMDA-treatment. There was a significant reduction in the total number of Iba1^+^ microglia/macrophages in *Rlbp1-CreERT:Ikkb^fl/fl^* retinas compared to *Rlbp1-CreERT* controls at 48hrs after NMDA damage **(Fig 3a-b)**. In addition to activation of resident microglia, reactive peripheral immune cells migrate into the retina after damage or disease^33–35^. Resident microglia and macrophages are CD45^(lo)^, whereas peripheral monocyte derived macrophages are CD45^(hi)^ expressing cells^36,37^. We identified CD45^+^/Iba1^−^ putative recruited monocyte-derived macrophages within the retina following NMDA-treatment; these cells likely originated from outside of the retina **(Fig 3a)**. We found a significant reduction in the total numbers of CD45^+^/Iba1^−^ immune cells in damaged retinas with *Ikkb-*cKO MG compared to controls **(Fig 3b)**. Taken together these data indicate that NFkB regulates MG-to-microglia communication and the recruitment of peripherally derived immune cells into damaged retinas.

**Figure 3:**
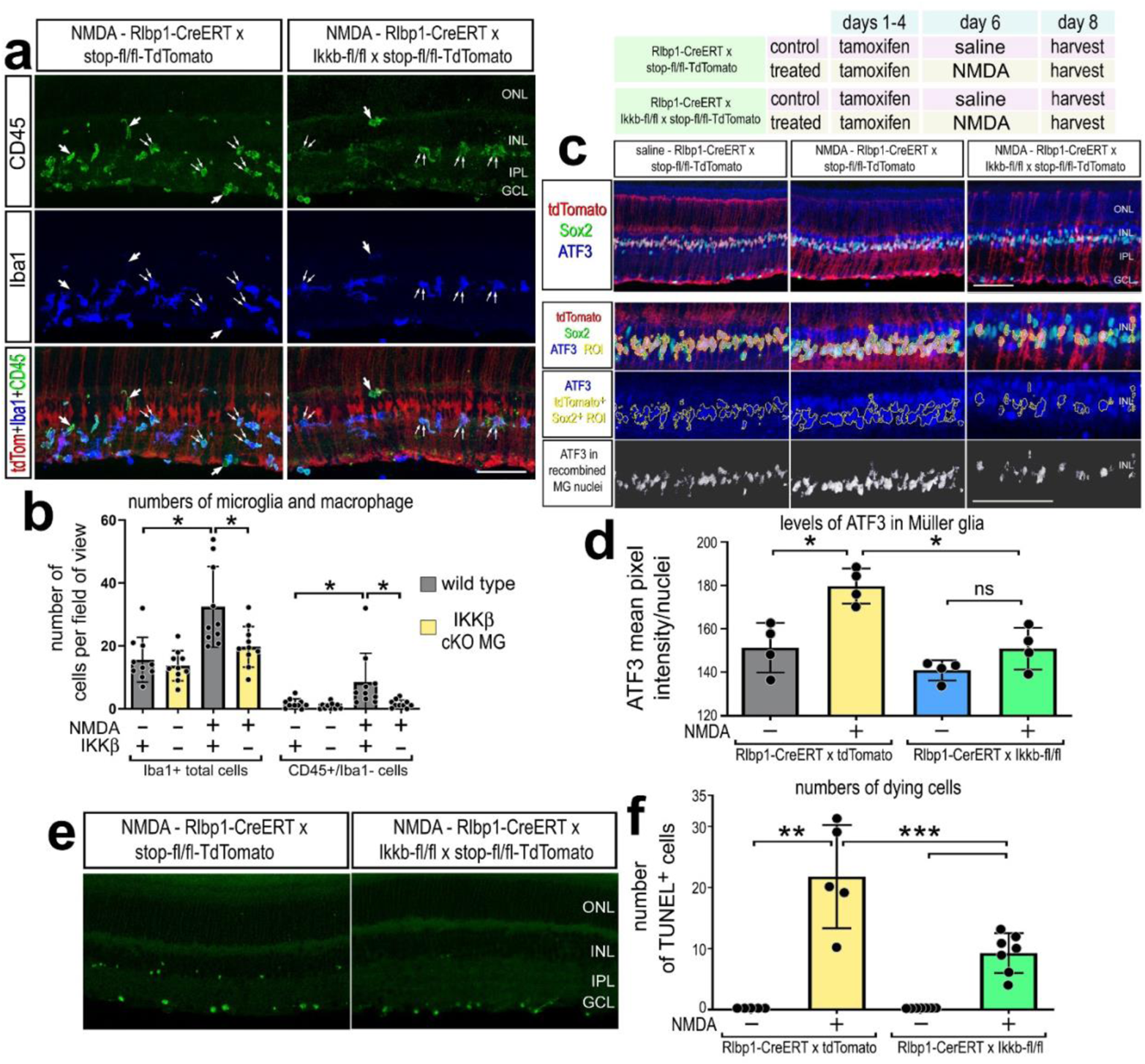
Conditional knockout of NFkB in Müller glia impairs immune cell responses after damage. *Rlbp1-creERT* and *Rlbp1-creERT*:*Ikkb*^fl/fl^ mice were injected IP with tamoxifen 1x daily for 4 consecutive days. Left eyes were injected with saline and right eyes were injected with NMDA on D6, and retinas were harvested on D8. **(a)** Retinal sections were labeled for CD45 (green), Iba1 (blue), and TdTomato (red). The histogram in **b** illustrates the mean number (±SD and individual data points) of Iba1^+^ or CD45^+^/Iba1^−^ cells. **(c)** Retinal sections were labeled for TdTomato (red), Sox2 (green), and ATF3 (blue, greyscale). The histogram in **d** illustrates the mean (±SD and individual data points) pixel intensity above threshold for ATF3 immunofluorescence within MG nuclei. **(e)** TUNEL assays were performed on retinal sections to identify dying cells. The histogram in **f** illustrates the mean number (±SD and individual data points) of TUNEL^+^ cells. Significance of difference (*p<0.05, **p<0.01) was determined by using one way ANOVA and Tukey’s post-hoc test.

We next probed for the expression of Activation Transcription Factors (ATFs) which are basic leucine zipper-containing cAMP-response element binding (CREB) proteins known to be involved in cellular stress responses, cell growth and survival, and are known to be downstream of NFkB-signaling^38–40^. Accordingly, we probed for levels of immunofluorescence for ATF3 in *Ikkb*-cKO MG. In undamaged retinas, ATF3-immunoreactivity was observed in the nuclei of MG and levels of immunofluorescence were significantly increased at 48hrs after NMDA-treatment **(Fig. 3c-d)**. There was a significant reduction in levels of ATF3 in the nuclei *Ikkb*-cKO MG after damage compared to levels seen in control MG in damaged retinas, and these levels were not significantly different from levels seen in the nuclei of control MG in undamaged retinas (Fig 3c-d).

Lastly, we assayed for changed in cell death considering previous studies indicating that inhibition of NFkB is neuroprotective against NMDA-induced damage in mammalian^16,17^ and avian^15^ retinas. We found a significant decrease in the number of TUNEL positive cells in retinas of *Rlbp1-CreERT:Ikkb^fl/fl^* mice compared to *Rlbp1-CreERT* control mice at 48hr after NMDA treatment **(Fig. 3e-f)**. Taken together, these data indicate that NFkB signaling is integrally involved in mediating the stress responses of MG and immune cells to neuronal damage and survival.

### scRNA-seq analyses of glial cells from retinas with *Ikkb*-cKO Müller glia

To further investigate the effects of disrupting NFkB signaling in MG we conducted scRNA-seq analysis to probe transcriptomic changes in retinal glia. *Rlbp1-CreERT* (control) mice and *Rlbp1-CreERT:Ikkb^fl/fl^* (*Ikkb-*cKO) mice received 4 consecutive daily IP doses of tamoxifen. Three days later both control and *Ikkb-*cKO mice received intravitreal injections of NMDA. Retinas were harvested for scRNA-seq 8 hours after NMDA treatment. Distinct types of retinal neurons and glia were identified based on expression of cell-distinguishing markers **(Supplemental Fig. 3**). UMAP ordering of MG revealed clusters of resting and activated MG of mixed library origin **(Fig. 4a,b)**. We identified differentially expressed genes (DEGs) between control MG and *Ikkb*-cKO MG, 603 DEGs were down-regulated, whereas there were 242 upregulated DEGs in *Ikkb*-cKO MG. Gene Ontology (GO) enrichment analysis of DEGs between control and *Ikkb*-cKO MG revealed significant downregulation of gene modules associated with the regulation of immune cell responses, inflammatory cytokine signaling, and signaling pathways known to influence NFkB-, Akt- and MAPK-signaling **(Fig. 4c)**. Additionally, chemokine signaling and leukocyte chemotaxis pathways were downregulated in *Ikkb*-cKO MG. Thus, we probed scRNA-seq libraries for *Ccl2* which is known to be involved in recruiting peripheral monocyte-derived cells into the retina^41–44^. We find *Ccl2* is expressed in endothelial cells, microglia, and is significantly upregulated in MG at 3, 6 and 12hrs after NMDA-treatment **(Supplemental Fig. 4j-k)**. We found a significant decrease in *Ccl2* expression levels in *Ikkb-*cKO MG compared to levels seen in WT MG **(Fig. 4d,f)**.

**Figure 4:**
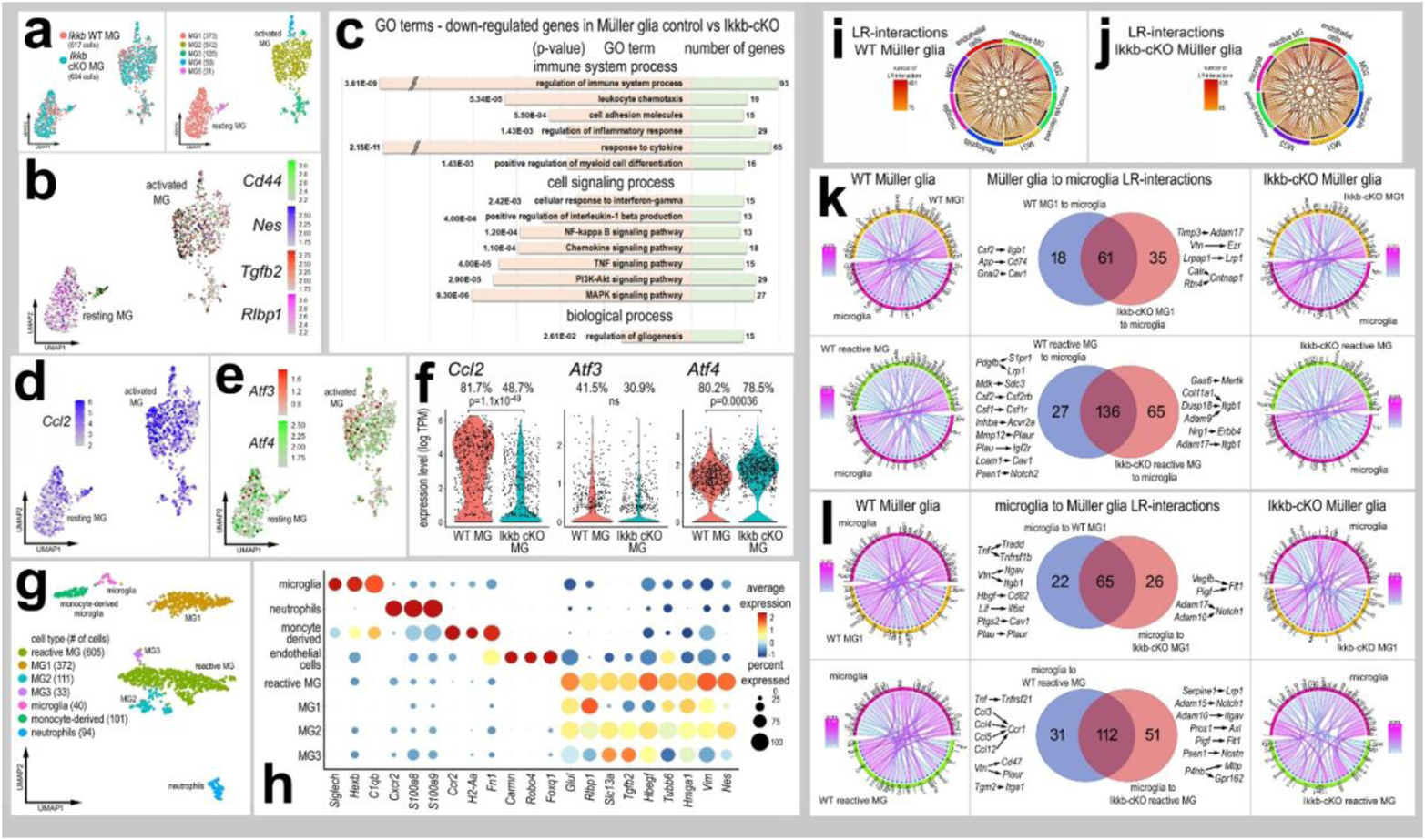
scRNA-seq analyses of glial cells from retinas with *Ikkb*-cKO Müller glia. UMAP plots of isolated and re-embedded MG from scRNA-seq libraries of *Rlbp1-creERT* (*Ikkb-*WT) and *Rlbp1-creERT:Ikkb*^fl/fl^ (*Ikkb-*cKO) retinas 8hr after NMDA **(a)**. Resting and reactive MG were identified by expression of *Rlbp1* or *Cd44, Nes*, and *Tgfb2* **(b)**. GO analysis on the significant (adjusted p-value ≤ 0.05) DEGs between *Ikkb*-WT and *Ikkb*-cKO MG **(c)**. UMAP plots **(d-e)** and violin plots **(f)** of MG clusters illustrate expression levels of *Ccl2, Atf3*, and *Atf4.* UMAP plots of aggregated MG and immune cells **(g)**. Dot plot distinct patterns of gene expression of various clusters and illustrates changes in resting and reactivity genes in MG clusters 1-3 **(h)**. Chord diagrams illustrate potential autocrine and paracrine ligand receptor (LR-) interactions generated from SingleCellSignalR for *Ikkb-*WT control and *Ikkb-*cKO libraries **(i-j)**. Chord plots show the 40 most significant LR-interactions from MG1 to microglia or from reactive MG to microglia from *Ikkb*-WT control and *Ikkb*-cKO libraries **(k)**. Chord plots show significant LR interactions from microglia to MG1 or to reactive MG from *Ikkb*-WT control or *Ikkb*-cKO libraries **(l)**.

We next probed for expression levels of ATFs which can be regulated by NFkB-signaling^40^. We found *Atf1* and *Atf2* are predominantly expressed by different types of retinal neurons, whereas *Atf3* and *Atf4* are widely expressed by retinal neurons and glia **(Supplemental Fig. 4d-e)**. *Atf3 and Atf4* are expressed at low levels in resting MG, but are upregulated at 3, 6 and 12hrs after damage **(Supplemental Fig. 4g**). At 8hrs after NMDA-treatment, the percentage of *Atf3* expressing MG was reduced by 10%, although levels of *Atf3* were not significantly different between control MG and *Ikkb*-cKO MG at this timepoint **(Fig. 4f)**. These findings are consistent with immunolabeling for ATF3 in MG at 48hrs after NMDA-treatment **(Fig. 3c,d)**.

Since deletion of *Ikkb* from MG resulted in diminished recruitment of reactive microglia/macrophage **(Fig. 3)**, we bioinformatically combined MG and immune cells and ordered these cells in UMAP plots **(Fig. 4g)**. The UMAP-clustered cells had distinct patterns of gene expression identifying clusters of microglia, neutrophils, monocyte-derived cells, endothelial cells, and 4 clusters of MG **(Fig. 4g-h)**. There was one cluster of highly reactive MG, characterized based on the expression of reactive glial genes *(Tgfb2, Vim, Hmgb1, Nes)*, while a separate cluster, MG1, expressed lower levels of genes associated with reactivity **(Fig. 4-h)**. We probed for changes in cell signaling networks and putative ligand-receptor (LR) interactions between immune cells and MG using SingleCellSignalR^45^. We focused our analyses on MG, microglia, and monocytes because there is significant evidence indicating signaling among these cells in the retina^46,47^. We identified 114 significant LR-interactions from MG1 signaling to microglia, of which 18 were specific to only control MG, and 35 were specific to *Ikkb*-cKO MG **(Fig. 4k)**. We identified 228 significant LR-interactions between reactive MG to microglia, with 27 specific to control MG and 65 specific to *Ikkb*-cKO MG **(Fig. 4k**). LR-interactions specific to control MG-microglia include *Csf1/2-Csf1r/Csf2rb*, which is involved in promoting macrophage differentiation and survival^29,48^. Additionally, LR-interactions that promote inflammation were identified in control libraries with WT MG (*App-Cd74, Inhba-Acvr2a, Mmp12-Plaur, Mdk-Sdc3*), and not in libraries with *Ikkb-*cKO MG **(Fig. 4k**).

We next assessed the significant LR-interactions from microglia to MG upon knockout of *Ikkb* in MG. We identified 117 significant interactions between microglia and MG, with 22 specific to libraries with WT MG and 26 specific to libraries with *Ikkb-*cKO MG **(Fig. 4l)**. We identified 194 significant LR-interactions between microglia and reactive MG, with 31 specific to libraries with WT MG and 51 specific to libraries with *Ikkb-*cKO MG **(Fig. 4l)**. LR-interactions enriched in *Ikkb*-cKO libraries include factors involved in modulating angiogenesis (*Vegfb/Pigf-Flt1*) as well as A Disintegrin and Metalloproteinase Domain (ADAM)-containing proteins to *Notch1* and *Itgav.* LR-interactions specific to control libraries are enriched for TNF to *Tradd/Tnfrsf1b/Tnfrsf21*, vitronectin to integrins/*CD47/Plaur*, and C-C motif chemokine ligands to *Ccr1* **(Fig. 4l)**. Taken together, these data indicate that NFkB signaling in MG is an essential signaling hub that mediates cross-talk between MG and microglia/macrophages and is involved in mediating glial reactivity and immune cell recruitment in response to damage.

### Inhibition of NFkB promotes Ascl1-mediated neuron regeneration

It has recently been shown that microglia provide signals to suppress the ability of MG to reprogram into neuronal cells following forced over expression of *Ascl1* in the mammalian retina^23^. Given our findings indicating that the ablation of microglia prevents the activation of NFkB in MG **(Fig. 2e-f)**, we hypothesized that actived NFkB in MG may suppress neurogenic reprogramming. To test this hypothesis we applied NFkB inhibitor (PGJ2), which potently suppresses NFkB activity **(Fig. 2c-d)**, to the retinas of *Glast-CreER:LNL-tTA:tetO-mAscl1-ires-GFP* mice^21^. To induce neuron regeneration from MG treatment with NMDA, to induce neuronal damage, and HDAC inhibitor trichostatin A (TSA) is required^21^. The paradigm of Ascl1+NMDA+TSA (hereafter referred to as ANT treatment) selectively drives expression of Ascl1 and GFP-reporter (to lineage trace) in MG to produce bipolar-like neurons and a few amacrine-like neurons that integrate into local circuitry and respond to light^21^. These regenerated neurons are formed primarily through transdifferentiation, but a subset of Ascl1-overexpressing MG proliferate prior to giving rise to new neurons^22^. We first validated the ANT paradigm to confirm efficacy and baseline numbers for neuronal regeneration. Consistent with previous results^21–23^, we found that treatment of retinas with NMDA and TSA significantly increased the percentage of GFP^+^/Otx2^+^ cells (bipolar-like cells) produced by Ascl1-overexpressing MG **(Supplemental Fig. 5c-d)**. Conversely, we found that treatment of retinas with NMDA and TSA significantly decreased the percentage of GFP^+^/Sox2^+^ cells (cells maintaining glial phenotype) in Ascl1-overexpressing MG **(Supplemental Fig.5a-b)**.

**Figure 5:**
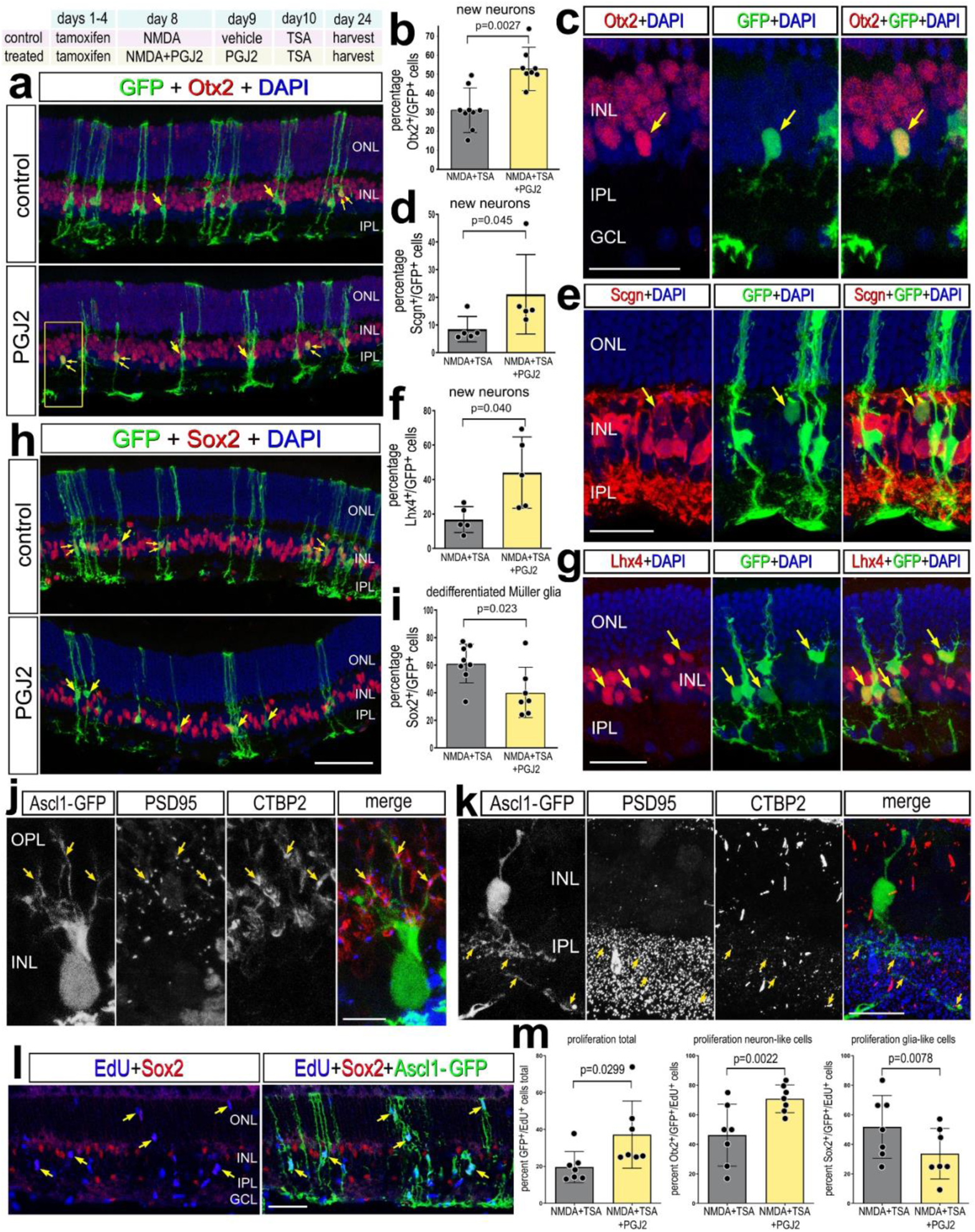
Inhibition of NFkB promotes Ascl1-mediated neuron regeneration. Experimental paradigm outlined at top. Tamoxifen was administered IP 1x daily for 4 consecutive days to *Glast*-*CreER:LNL-tTA:tetO-mAscl1-ires-GFP* mice. NMDA was injected intravitreally in left (control) eyes and NMDA+ PGJ2 in right (treated) eyes on D8, vehicle ± PGJ2 on D9, TSA ± PGJ2 on D10, and retinas were harvested 2 weeks after TSA injection. Retinal sections were labeled for GFP (green), DAPI (blue), and Otx2 (red; **a,c**), Scgn (red; **e**), Lhx4 (red; **g**), or Sox2 (red; **h**). Arrows in **a,c,e,h** and **g** indicate cells double-labeled for GFP and Otx2, Scgn, Lhx4 or Sox2, and small double-arrows indicate GFP alone. Histogram illustrate the mean percentage (±SD and individual data points) of GFP^+^ cells that are Otx2^+^ **(b)**, SCGN^+^ **(d)**, Lhx4^+^ **(f)**, or Sox2^+^ **(i)**. Significance of difference (p-values shown) was determined by using a paired t-test. Retinal sections were labeled for GFP, PSD95, and CTBP2 to identify apposing pre- and post-synaptic markers in the OPL **(j)** and IPL **(k)**. **(l-m**) EdU was administered in the drinking water 24h prior to damage and sustained until harvesting retinas 4 days after TSA treatment. Retinal sections were labeled for GFP (green), Sox2 (red), EdU (blue) **(I**). Single arrows in **l** indicate GFP^+^/EdU^+^ cells. The histogram in **(m**) illustrates the mean percentage (±SD and individual data points) of GFP^+^ cells that are EdU^+^, and the percent of EdU^+^/GFP^+^ cells that are Otx2^+^ or Sox2^+^**(m**). The calibration bars represent 50 µm. Abbreviations: ONL, outer nuclear layer; INL, inner nuclear layer; IPL, inner plexiform layer; GCL, ganglion cell layer.

We next tested whether inhibition of NFkB increased the formation of neurons derived from Ascl1-overexpressing MG. Ascl1 expression in MG was activated by IP delivery of 4 consecutive daily doses of tamoxifen in adult mice (P90-P140). This was followed by intravitreal injection of NMDA or NMDA+PGJ2 on D8 and injection of TSA+vehicle or TSA+PGJ2 on D10 **(Fig. 5)**. Eyes were harvested and retinas were processed for immunolabeling 2 weeks after the final injection. We found a significant increase in the proportion of Ascl1-overexpressing GFP^+^ cells that were co-labeled for Otx2, Scgn, or Lhx4 (but not HuC/D) in retinas treated with NFkB inhibitor **(Fig 5a-g)**, suggesting increased formation of bipolar-like neurons. Conversely, we found a significant decrease in the proportion of GFP^+^/Sox2^+^ MG **(Fig 5h-i)**, suggesting fewer Ascl1-MG remaining as glia. Additionally, GFP^+^ MG-derived neurons had morphologies distinct from mature MG, with spherical rather than fusiform cell bodies and processes retracted from the outer limiting membrane **(Fig. 5c,e,g**). Cells with neuronal morphology may have formed synapses with photoreceptors and inner retinal neurons indicated by apposition of synaptic markers CtBP2 with Psd95 on GFP^+^ processes in the OPL and IPL **(Fig. 5j,k)**.

ANT-treatment is known to stimulate some Ascl1-overexpressing MG to re-enter the cell cycle before generating neuron-like cells^22^. Thus, we probed for proliferation in ANT+PGJ2 treated retinas to determine whether inhibition of NFkB promoted the proliferation of Ascl1-overexpressing MG. To identify proliferating cells we applied EdU in the drinking water prior to damage and sustained the EdU exposure until harvesting retinas at 4 days after TSA treatment. We found a significant increase in the proportion of EdU^+^/GFP ^+^ cells in ANT+PGJ2 treated retinas compared to ANT alone **(Fig. 5l-m)**. This finding suggests that inhibition of NFkB promotes proliferation of Ascl1-overexpressing MG. Some of these EdU^+^/GFP ^+^ cells were positive for Sox2, while some EdU^+^/GFP ^+^ were negative for Sox2 **(Supplemental Fig. 6)**. There was a significant increase in the proportion of EdU^+^ /Ascl1-GFP ^+^ MG that were positive for Otx2 in ANT+PGJ2 treated retinas compared to controls, and a significant decrease in the proportion of EdU^+^ /Ascl1-GFP ^+^ MG that were positive for Sox2 in ANT+PGJ2 treated eyes compared to controls **(Fig. 5l-m, Supplemental Fig. 6)**. This suggests that inhibition of NFkB stimulates proliferation of cells with bias toward acquiring neuronal phenotype.

**Figure 6:**
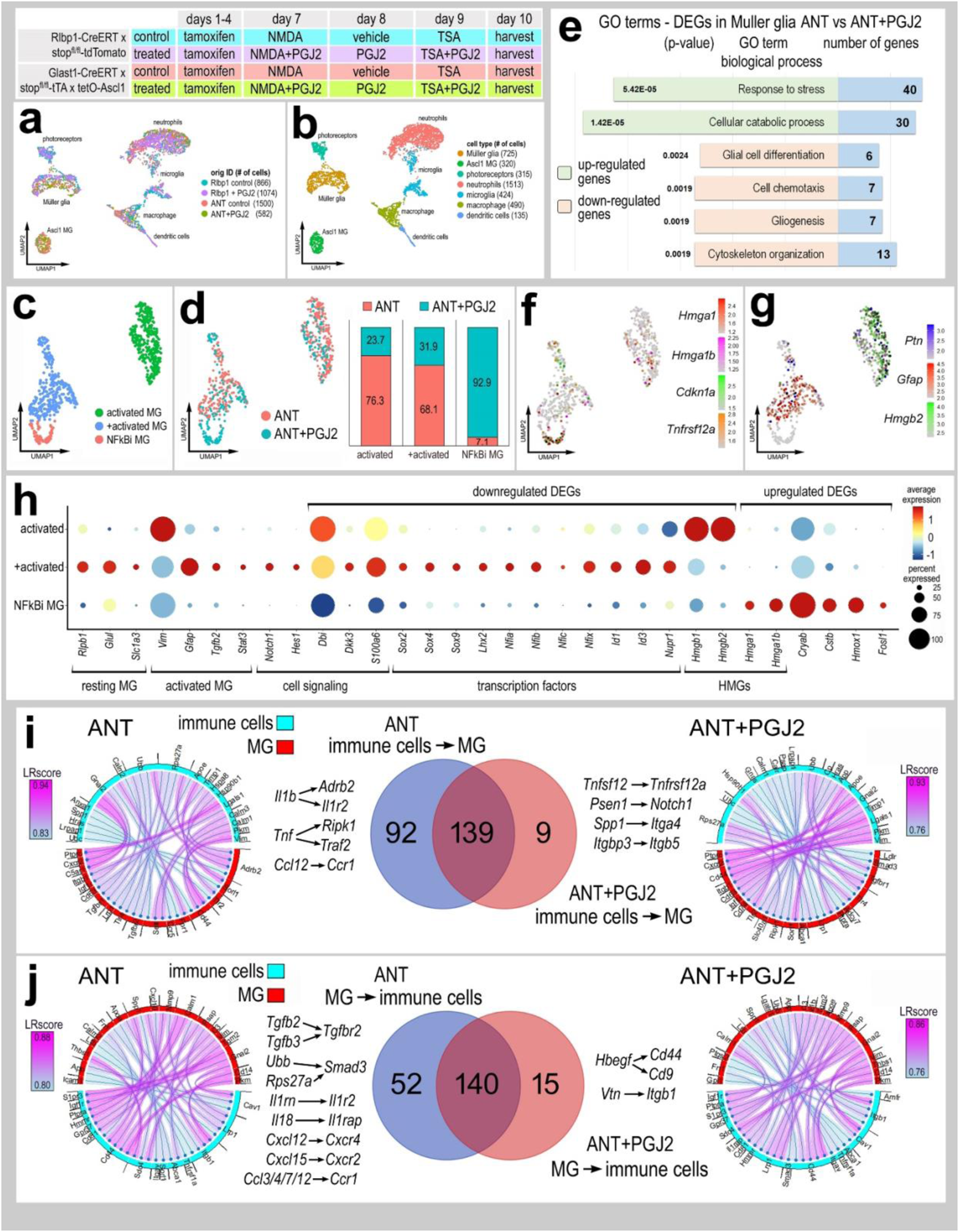
Inhibition of NFkB signaling and changes in transcriptomic profiles. Experimental paradigm outlined at top. scRNA-seq libraries were prepared from *Rlbp1-CreERT* and *Glast*-*CreER:LNL-tTA:tetO-mAscl1-ires-GFP* mice. Adult mice were treated with tamoxifen IP 1x daily for 4 consecutive days. NMDA was intravitreally injected into left (control) eyes and NMDA+PGJ2 in right (treated eyes) on D8, vehicle ± PGJ2 on D9, TSA ± PGJ2 on D10, and retina were harvested 24h after TSA treatment. Cells were FACS enriched for either GFP^+^ and CD45^+^/Cd11b^+^ (from Ascl1 over-expressing mice) or TdTomato^+^ cells and CD45^+^/Cd11b^+^ cells (from *Rlbp1-CreERT* mice). UMAP plots illustrate aggregated scRNA-seq libraries for sorted MG and immune cells **(a)**. Clusters of different types of cells were identified **(b)**. UMAP plots of isolated and re-embedded MG from scRNA-seq libraries of ANT (Ascl1-NMDA-TSA) and ANT+PGJ2 treated retinas **(c-d)**. Stacked bar histograms represent occupancy of each cluster by treatment (ANT or ANT+ PGJ2) **(d)**. GO analysis of significant (adjusted P value ≤.05) DEGs in MG from ANT and ANT+PGJ2 libraries **(e)**. UMAP plots of MG clusters illustrate patterns of expression of *Hmga1, Hmga1b, Cdkn1a, Tnfsf12a* **(f)** or *Ptn, Gfap, Hmgb2* **(g)**. Changes in resting- and reactivity-associated genes in MG clusters 1-3 were illustrated in dot plot, with dot size representing percentage of cells expressing and dot color representing level of expression **(h)**. Ligand receptor (LR-) interactions were assessed by using SingleCellSignalR. Chord plots illustrate the 40 most significant LR-interactions from immune cells to MG **(i)** or from MG to immune cells **(j)** from ANT (control) and ANT+PGJ2 (treated) conditions. The Venn diagrams illustrate numbers of common and differential LR-interactions between ANT (control) and ANT+PGJ2 (treated) conditions.

Additionally, we probed for changes in cell death since previous studies have shown that inhibition of NFkB is protective against excitotoxic retinal damage^16,17^, and increased levels of damage from the drug treatment could result in increased proliferation. However, we found a significant decrease in the number of TUNEL-positive cells at 24hrs after NMDA-treatment in ANT retinas treated with PGJ2 compared to ANT controls **(Supplemental Fig. 6)**. Taken together, these data indicate the inhibition of NFkB is neuroprotective and promotes reprogramming of Ascl1-overexpressing MG into bipolar-like neurons.

### Inhibition of NFkB and changes in transcriptomic profiles

We next sought to identify gene regulatory networks (GRNs) that are changed in MG upon inhibition of NFkB. We performed the ANT±PGJ2 regeneration paradigm on both *Glast-CreER:LNL-tTA:tetO-mAscl1-ires-GFP* mice and on *Rlbp1-CreERT* mice as controls. We performed FACs to enrich for GFP^+^ Ascl1-overexpressing MG, or TdTomato^+^ control MG, as well as CD45^+^/Cd11b^+^ microglia and macrophages. We generated scRNA-seq libraries for sorted glial cells from the following transgenic lines and treatment conditions: (1) *Rlbp1-CreERT+*NMDA+TSA, (2) *Rlbp1-CreERT+*NMDA+TSA+PGJ2, (3) ANT, and (4) ANT+PGJ2 **(Fig. 6a)**. We identified distinct clusters of cells that were identified as microglia, monocyte derived macrophages, neutrophils, dendritic cells and MG, as well as some photoreceptor cells from contamination **(Fig. 6b, Supplemental Fig. 7)**.

**Figure 7:**
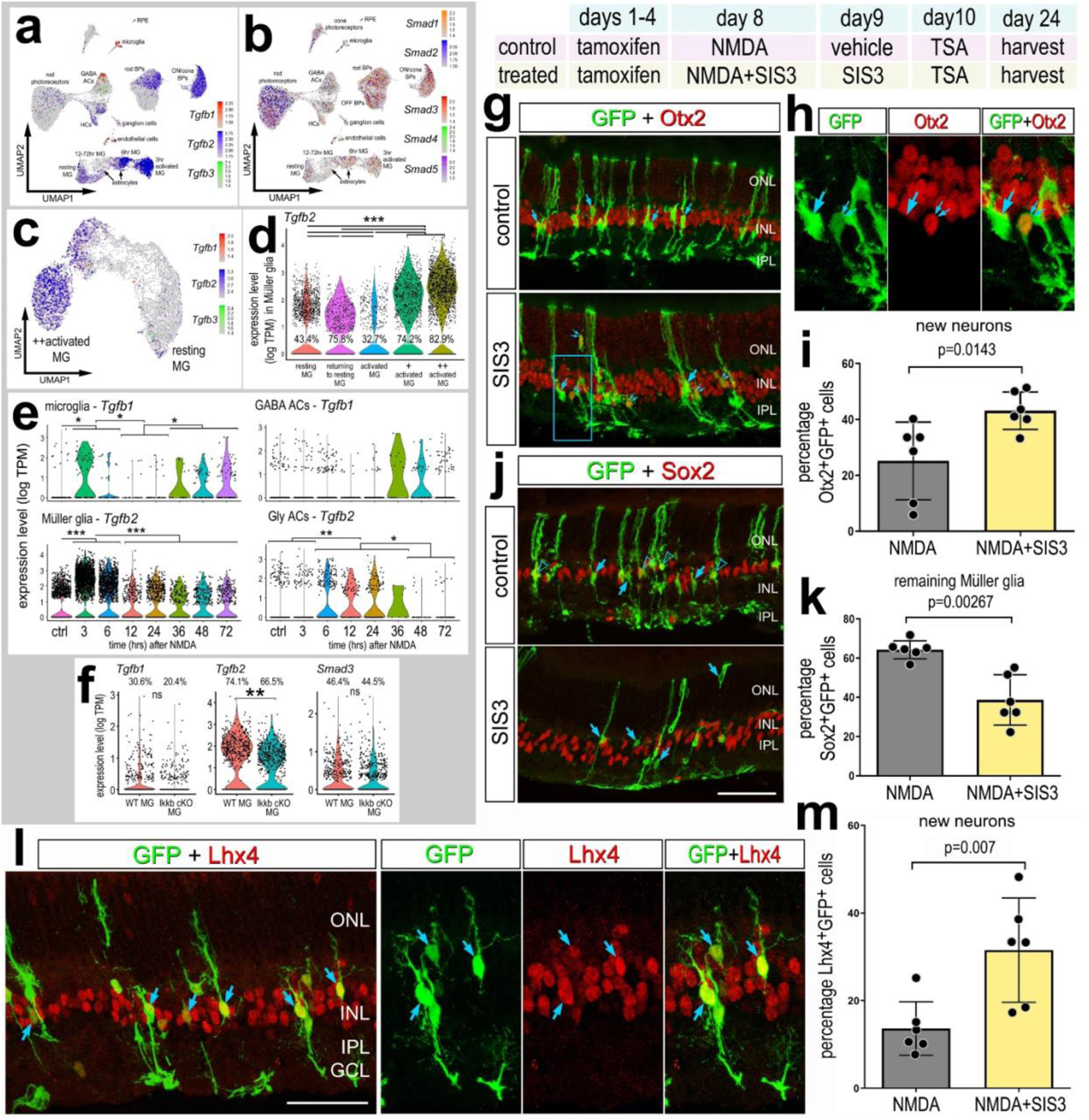
Inhibition of TGFb/Smad3 promotes neurogenesis. Aggregated scRNA-seq libraries were prepared from wild type control retinas and retinas 3, 6,12, 24, 36, 48 and 72hr after NMDA damage, and numbers of cells per library or per cluster in parentheses (see Fig.1 for legend). UMAP plots illustrate expression patterns of *Tgfb1, Tgfb2, Tgfb3* **(a**) or *Smad1, Smad2, Smad3, Smad4, Smad5* **(b**). UMAP plots of isolated and re-embedded MG illustrate patterns expression of *Tgfb1, Tgfb2, and Tgfb3* across a gradient of resting and activated MG from different times after NMDA treatment **(c)**. Violin plots illustrate expression levels and percentage of expressing for *Tgfb2* in MG **(d)**, and *Tgfb1* and *Tgfb2* in microglia, GABAergic amacrine cells, glycinergic amacrine cells and MG **(e)**. Violin plots in **(f)** illustrated expression levels of *Tgfb1/2* and *Smad3* from *Ikkb*-WT control MG and *Ikkb*-cKO MG **(f)**. Significance of difference (**p<0.01, ***p<0.001) was determined by using a Wilcoxon rank sum with Bonferroni correction **(d-f)**. Paradigm: tamoxifen was administered IP 1x daily for 4 consecutive days in *Glast*-*CreER:LNL-tTA:tetO-mAscl1-ires-GFP* mice, intravitreal injection of NMDA±SIS3 on D8, TSA ± SIS3 on D9, and retinas were harvested 2 weeks after the last injection **(g-m)**. Retinal sections were labeled for GFP (green), Otx2 (red; **g-h**), Sox2 (red; **j**), or Lhx4 (red; **l**). Histograms illustrate the percentage (±SD and individual data points) of GFP^+^ cells that are Otx2^+^ **(i)**, Sox2^+^ **(k)**, or Lhx4^+^ **(m)**. Significance of difference (p-values shown) was determined by using a paired t-test. Arrows in **g-h** indicate GFP^+^ cells negative for Otx2, small double-arrows in **g-h** indicate GFP^+^/Otx2^+^ cells. Arrows in **j** indicate GFP^+^ cells negative for Sox2, and hollow arrow-heads indicate GFP^+^/Sox2^+^ cells. Arrows in **l** indicate GFP^+^/Lhx4^+^ double-labeled cells. Calibration bars represent 50 µm. Abbreviations: ONL, outer nuclear layer; INL, inner nuclear layer; IPL, inner plexiform layer; GCL, ganglion cell layer.

To probe for changes in gene expression specifically in MG we isolated and re-normalized MG from all 4 libraries. UMAP plots revealed 2 clusters of Ascl1-overexpressing MG containing cells from both ANT (control) and ANT+PGJ2 (treated) libraries (Ascl1 MG1 and Ascl1 MG2); the cells were distinguished by differential expression of genes for resting glia (*Rlbp1, Id1, Id3* and *Nfix)* or activated glia (*Ptn, Hmgb1* and *Hmgb2*) **(Supplemental Fig. 7d-e)**. We also identified a cluster of cells containing both *Rlbp1-CreERT* control (NMDA) and *Rlbp1-CreERT* treated (NMDA+PGJ2) cells. This cluster of cells maintained high expression of resting glial genes such as *Rlbp1, Slc1a3, Dkk3* and *Glul* **(Supplemental Fig. 7d-e)**. Finally, we identified one cluster of cells consisting of only NFkB-inhibitor (NFkBi) treated cells (*Rlbp1-CreERT* treated (NMDA+PGJ2) and ANT+PGJ2) **(Supplemental Fig. 7d)**. This cluster of cells had a significant reduction in expression of pro-glial genes including *Lhx2, NFia/b/x, Hes1* and *Id1/3* **(Supplemental Fig. 7e)** ^13,49,50^. Additionally, the NFkBi cluster had decreased expression of genes associated with activation of glial reactivity including *Stat3, Notch2*, and *Tgfb2* **(Supplemental Fig. 7e)**.

We next isolated the MG from ANT (control) and ANT+PGJ2 (treated) libraries to probe for differences in Ascl1-overexpressing MG resulting from NFkB inhibition **(Fig. 6c-d)**. We performed GO analysis on significant up- or downregulated DEGs between control and treated libraries. GO enrichment analysis revealed upregulated groups of genes associated with cell stress and catabolism in Ascl1-MG treated with NFkB inhibitor **(Fig. 6e)**. GO enrichment analysis revealed downregulated groups of genes associated with glial differentiation and development, cytoskeleton organization, and chemotaxis in Ascl1-MG treated with NFkB inhibitor **(Fig. 6e)**. UMAP plots of MG from control and treated libraries revealed one cluster of exclusively ANT+PGJ2 cells (NFkBi MG cluster), and two clusters with mixed origin identity **(Fig. 6c-d)**. The NFkBi MG cluster showed enriched expression of *Hmga1, Hmga1b, Cdkn1a*, and *Tnfrsf12a* **(Fig. 6f)**. *Hmga1* is a high mobility group AT-hook 1 gene, which is an important factor driving the transition from reactive MG to progenitor state in zebrafish^13^. *Hmga1* expression is induced in reactive MG in the zebrafish retina after damage, and knockdown of *Hmga1* decreased MG proliferation and resulted in increased MG reactivity^13^. Thus, the increased *Hmga1* expression is a promising candidate underlying the increase in transition of reactive glia to a progenitor state in the mammalian retina following inhibition of NFkB.

The NFkBi MG cluster had a significant reduction in expression of reactivity genes *(Gfap, Ptn, Tgfb2, Hmgb1/2, Sox9)* **(Fig. 6g-h)**. Additionally, there was a significant reduction in expression of pro-glial factors that suppress neurogenesis, including *Nfia/b/x*^13,49^, *Id3 and Hes1*^51^ **(Fig. 6h**). Taken together, these data indicate that inhibition of NFkB decreases expression of genes associated with glial reactivity and diminishes pro-glial/anti-neuronal transcriptional networks.

We used SingleCellSignalR^45^ to identify putative Ligand-Receptor (LR) interactions among DEGs in immune cells (microglia, macrophages, neutrophils) and MG in ANT control and ANT+PGJ2 conditions **(Fig. 6i-j)**. Significant LR-interactions from immune cells to MG included *Il1b-IL1r2, TNF-Traf*, and *Ccl12-Ccr1* in ANT control conditions but are absent in ANT+PGJ2 conditions **(Fig. 6i)**. Comparatively, *Tnfsf12-tnfrsf12a, Spp1-Itga4*, and *Itgbp3-Itgb5* were enriched in ANT+PGJ2 libraries **(Fig. 6i)**. Similarly, significant LR-interactions from MG to immune cells included *Tgfb2/3-Tgfbr2, Ubb-Smad3, Il18-Il1r2, Cxcl12/15-Cxcr4/2, Ccl3/4/7/12-Ccr1* in ANT control conditions, but are absent in ANT+PGJ2 treated conditions **(Fig. 6j)**. Comparatively, *Hbegf-Cd44/Cd9* and *Vtn-Itgb1* LR-interactions were significantly enriched in ANT+PGJ2 conditions **(Fig. 6j)**. These findings suggest that NFkB signaling in MG facilitates pro-inflammatory cascades, recruitment of peripheral immune cells into the retina, and activation of pro-glial signals and transcriptional networks coordinated through communication with immune cells.

### Inhibition of TGFb/Smad3 promotes neurogenesis

TGFb/Smad signaling may be a point of divergence regarding the responses of MG to damage across species^5,12,52^. In the fish retina, TGFb/Smad3 signaling is activated MG in response to light damage, but is then rapidly repressed^53^, and inhibition of TGFb signaling promotes retinal regeneration in the fish retina^54^. Tgfb1/2 promotes reactivity/gliosis in MG in the mouse retina, while Tgfb3 promotes MG-mediated regeneration in the zebrafish retina^55^. In the chick retina, TGFb/Smad2/3 signaling suppresses the formation of MGPCs and acts in opposition to BMP/Smad1/5/8 signaling which promotes the formation of MGPCs^56^. We found a significant reduction in expression of *Tgfb2* in MG upon inhibition of NFkB with ANT+PGJ2 treatment **(Fig. 6h)** or *IKKb*-cKO in MG **(Fig. 7f)**. Thus, we tested whether inhibition of TGFb/Smad signaling alone influenced *Ascl1*-mediated reprogramming of MG. First, we analyzed expression patterns of TGFb ligands and Smad signal transducers in scRNA-seq libraries of WT NMDA damaged mouse retina **(Fig. 7a-e)**. We found that *Tgfb2 is* highly expressed in activated MG and in MG that are returning to a resting state **(Fig. 7c-e)**. *Tgfb3* is not widely expressed **(Fig. 7a,c)**. Additionally, *Tgfb1/2* are expressed by microglia, amacrine cells, endothelial cells, rod bipolar cells and ON/cone bipolar cells **(Fig. 7e)**. Levels of *Tgfb1* in microglia and *Tgfb2* in glycinergic amacrine cells significantly changed after damage **(Fig. 7b,e**). *Smads1-5* were expressed by scattered cells in nearly all types of neurons and glia **(Fig. 7b)**.

Next, we applied SIS3, a small molecule known to selectively inhibit Smad3^57^, to the ANT regeneration paradigm. Previous studies have shown that treatment with SIS3 increases the formation of proliferating MGPCs in damaged chick retinas^56^. The ANT+SIS3 treatment induced a significant increase in the proportion of GFP^+^/Otx2^+^ **(Fig. 7g-i)** and GFP^+^/Lhx4^+^ MG-derived neurons **(Fig. 7l-m)**. By contrast, ANT+SIS3 treatment induced a significant decrease in the proportion of GFP^+^/Sox2^+^ MG that remained as glia **(Fig. 7j-k)**. Taken together, these data indicate that NFkB coordinates with TGFb2/Smad3 signaling to suppress the neurogenic potential of MG.

### Inhibition of Id transcription factors promotes neurogenesis

Our findings suggest that levels of *Id1* and *Id3* are significantly reduced in Ascl1-overexpressing MG by NFkB inhibition **(Fig. 8f)** and upon *Ikkb*-cKO in MG **(Fig. 8g**). Ids are pro-glial transcriptional regulators that can bind to and prevent pro-neural bHLH transcription factors from binding to DNA^58^, and repress neurogenesis^51,59^. Ascl1-overexpressing MG that remain as glia and don’t differentiate into neurons express higher levels of *Id1* and *Id3*^22^. Analysis of our scRNA-seq databases indicated significant and rapid downregulation of *Id1* (3hrs) and upregulation of *Id2* and *Id3* (3-6hrs) in MG after NMDA-treatment, whereas *Id4* was not expressed at significant levels in control and treated MG **(Fig. 8a-e**). Additionally, levels of *Id2* and *Id3* are significantly reduced in MG upon *Ikkb*-cKO or inhibition of NFkB **(Fig. 8f)**.

**Figure 8:**
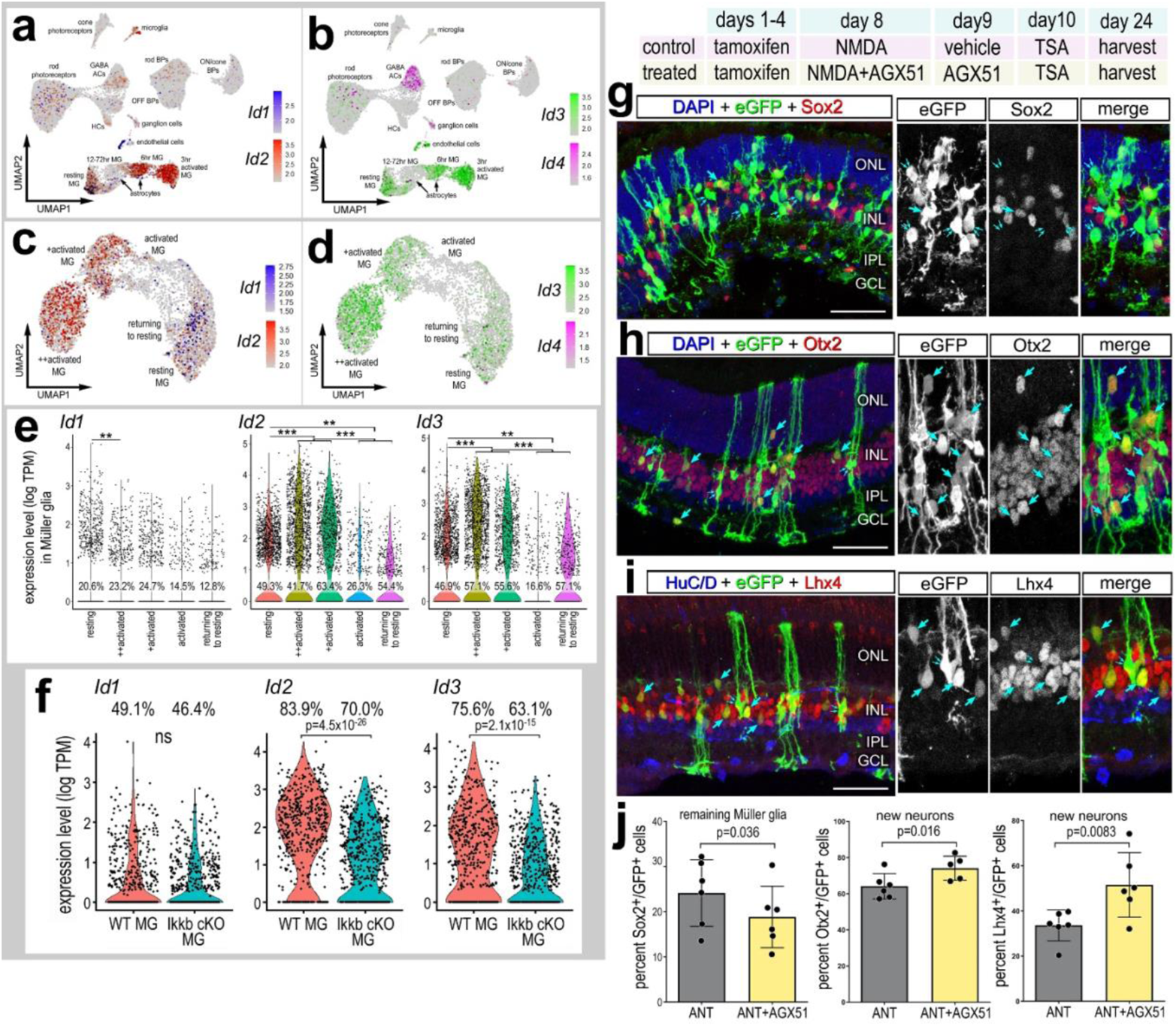
Inhibition of ID transcription factors promotes neurogenesis. Aggregated scRNA-seq libraries were prepared from wild type control retinas and retinas 3, 6,12, 24, 36, 48 and 72hr after NMDA damage. Numbers of cells per library or numbers of cells per cluster are listed in parentheses. UMAP plots illustrate expression of *Id1, Id2* **(a**) or *Id3, Id4* **(b**). UMAP plots of isolated and re-embedded WT MG illustrate expression of *Id1, Id2* **(c**) or *Id3, Id4* **(d**). Violin plots for isolated MG show expression of *Id1, Id2, Id3* from WT retinas **(e)** UMAP and violin plots of isolated and re-embedded ANT±PGJ2 MG show expression of *Id1* and *Id3* **(f)** or from *Ikkb-*WT control and *Ikkb-* cKO libraries **(g)**. Significance of difference (**P<0.01, ***P<0.001) was determined by using a Wilcoxon rank sum with Bonferroni correction. Experimental paradigm outlined: tamoxifen administered IP 1x daily for 4 consecutive days in *Glast-CreER:LNL-tTA:tetO-mAscl1-ires-GFP* mice, NMDA ± AGX51 on D8, vehicle ± AGX51 on D9, TSA ± AGX51 on D10, and retinas were harvested 3 weeks after TSA **(h-k)**. Representative images of treated eyes, retinal sections were labeled for GFP (green), Sox2 (red; **h**), Otx2 (red; **i**), or Lhx4 (red; **j**). Single arrows in **h** indicate GFP^+^/Sox2^+^ cells, double arrows indicate GFP^+^/Sox2^−^ cells. Single arrows in **(i)** and **(j)** indicate GFP^+^/Otx2^+^ or GFP^+^/Lhx2^+^ cells, respectively. Histograms in illustrate the percentage (±SD and individual data points) of GFP^+^ cells that are Sox2^+^, Otx2^+^, or Lhx4^+^ **(k)**. Significance of difference (p-values shown) was determined by paired t-test. Calibration bars in images represent 50 µm. Abbreviations: ONL, outer nuclear layer; INL, inner nuclear layer; IPL, inner plexiform layer; GCL, ganglion cell layer.

Thus, we next investigated whether inhibition of Ids was sufficient to promote increased neurogenesis from *Ascl1*-overexpressing MG. We applied a small molecule Id inhibitor, AGX51, to the ANT paradigm. AGX51 inhibits the interactions with Id1-E47 and targets Id1, Id2, Id3, and Id4 for ubiquitination and degradation^60^. We find a significant increase in the proportion of GFP^+^/Otx2^+^ **(Fig. 8i,k)** and GFP^+^/Lhx4^+^ **(Fig. 8j-k)** MG-derived neurons in ANT retinas treated with Id inhibitor. We found a corresponding decrease in the proportion of GFP^+^/Sox2^+^ MG that de-differentiated in response to Id inhibitor **(Fig. 8h,k)**. Taken together, these data indicate that NFkB signaling is involved in promoting expression of Ids, which in turn may repress neurogenesis of Ascl1-overexpressing MG.

## Discussion

The process of MG-mediated retinal regeneration is contingent upon a balance between MG activation, reactive gliosis, de-differentiation and neurogenesis. MG reactivity is necessary prior to proceeding to neurogenic reprogramming in lower vertebrates^13,61^. However, in mammalian MG, reactivity and quiescence are tightly regulated by different networks of genes^13^. Our findings suggest that NFkB drives MG reactivity and pro-glial/quiescence programs, thereby blocking transition toward a neurogenic progenitor-like state, and this occurs rapidly within 2 days after damage. By comparison, the differentiation of neuron-like cells from *Ascl1*-overexpressing MG is a slow process that takes 2-3 weeks after retinal damage in adult rodents^21^. Our findings indicate that there is activation of NFkB-signaling within 48hrs of damage, and inhibition of NFkB during this early time increases numbers of neuron-like cells that differentiate over the following 2 weeks. Similarly, inhibition of TGFb-signaling or Id transcription factors during the first 2 days after damage is sufficient to increase numbers of neuron-like cells derived from *Ascl1-*overexpressing MG. These findings indicate that shortly after acute retinal injury, MG activate a program involving NFkB signaling, TGFb signaling, and Id transcription factors that suppress neurogenic potential, and likely act to restore and maintain glial phenotype. This is consistent with our findings in the chick retina that activation of NFkB signaling suppresses the formation of MGPCs and promotes differentiation of progeny into glial cells^15^.

NFkB signaling is a well-established mediator of inflammation following damage or stress, in addition to regulating genes involved in cell survival, apoptosis, proliferation and differentiation^27^. NFkB signaling is activated by a variety of cytokines, primarily members of the interleukin (IL) and TNF ligand/receptor families^62^. NMDA-induced excitotoxic damage in the retina is known to activate NFkB in MG^17^. Here we establish an essential role of microglia in activating NFkB within MG in response to damage. Microglia are the resident immune cells of the retina and are a primary source of pro-inflammatory cytokines. In response to retinal injury, microglia acquire an activated phenotype, associated with inflammation, and migrate to the site of injury^63^. There is evidence showing bi-directional communication between microglia and MG that regulates inflammatory responses after retinal injury^47^. In NMDA-damaged retinas, reactive microglia appear to be the only source of *Il1a, Il1b* and *Tnf* ^28^, pro-inflammatory cytokines that likely activate NFkB signaling. Our data establishes NFkB signaling as a major hub of communication from microglia to MG that is likely activated by cytokines.

Our data indicate that activation of NFkB in MG induces an inflammatory response in MG that involves activation of chemokine pathways involved in recruiting peripheral immune cells into the retina. Signaling through Ccl2 and Ccr2 is essential for infiltration of monocytes into the retina^42,64^. We observed a significant reduction in expression of *Ccl2* in MG when NFkB signaling had been conditionally knocked-out.

This reduction of *Ccl2* in MG may be responsible for the diminished accumulation of reactive microglia and recruitment of peripheral immune cells to retinas where NFkB signaling was disrupted in MG. Consistent with prior studies suggesting that inhibition of NFkB in the retina is neuroprotective^16,17^, we found that selective deletion of *Ikkb* from MG resulted in neuroprotection. These findings are consistent with observations that factors, such as CNTF or FGF2, that activate cell signaling primarily in MG provide potent neuroprotection^65,66^, implicating MG as key mediators of cell survival in the retina.

Reactive microglia and macrophages are known to influence the reprogramming response of MG after damage. In the zebrafish, inflammatory signals are necessary for MG reprogramming and ablation of microglia in the retina prior to injury impairs the process of neuronal regeneration^33,67^. In the chick retina, the ablation of microglia in the retina prevents the formation of proliferating MGPCs^68^, but this can be rescued by delivery of a pro-inflammatory cytokine (TNFSF15) or by pharmacological activation of NFkB-signaling^15^. However, in mammalian retinas, microglia interfere with MG-mediated neuronal regeneration; ablation of microglia promotes Ascl1-mediated neurogenesis and treatment with TNF decreases Ascl1-mediated MG reprogramming^23^. These findings are consistent with our results indicating that activation of NFkB in MG is mediated by reactive microglia, and inhibition of NFkB promotes the formation of MG-derived neurons following overexpression of Ascl1.

Following retinal damage, zebrafish MG activate gene networks that drive reactivity prior to de-differentiation and entering the cell cycle as a progenitor cell^13^. Similarly in chick retinas, MG upregulate reactivity genes and proceed to de-differentiate and become proliferating progenitor cells or re-enter quiescence^13^. However, MG in damaged mouse retinas upregulate gene networks associated with reactivity and networks that rapidly restore quiescence^13^.The networks driving restoration of quiescence in mammalian MG include Nfia/b/x, Hes5, and Ids, and Nfia/b/x knockout is sufficient to promote neuronal reprogramming of MG in damaged mouse retinas^13^. We find that inhibition of NFkB in Ascl1-overexpressing MG decreases expression of genes (Nfia/b/x, Hes5, Ids) that promote pro-glial and quiescence networks. We propose that decreased levels of genes associated with pro-glial and quiescence networks underlie the increased neurogenesis following NFkB inhibition. In addition, inhibition of NFkB decreases expression of genes associated with reactivity networks involving *Tgfb2, Stat3*, and *Notch1/Hes*. This is consistent with a recent study demonstrating that the inhibition of Jak/Stat-signaling results in increased neurogenesis from Ascl1-overexpressing MG^22^.

The increased neurogenesis from Ascl1-overexpressing MG treated with NFkB inhibitor may be mediated by HMG factors including *Hmga1* and *Hmga1b*. Hmga1 is a non-histone chromatin modifying protein^69^ and is highly expressed in malignant cancers and stem cells during embryogenesis^70,71^. Hmga1 is expressed during embryonic development, but is undetectable in most adult tissues, except for cells residing in adult stem cell niches where this factor plays an important role in promoting stemness^72^.

Further, Hmga1 enhances reprogramming of somatic cells into induced pluripotent stem cells when combined with Sox2, Oct4, c-Myc and Klf4^73^. Hmga1 expression is induced in reactive MG in the zebrafish retina after damage and is required for the transition from reactivity to a progenitor-like state; knockdown of Hmga1 decreased proliferation of MGPCs and increased MG reactivity^13^. We find a significant increase in expression of *Hmga1* and *Hmga1b* following NFkB inhibition. Thus, we propose that *Hmga1* and *Hmga1b* may enhance the transition from a reactive MG state to progenitor-like state in the mammalian retina.

In addition to its role as a master regulator of cellular stress, ATF3 has been established to promote pro-glial gene networks and is a driver of astrocyte differentiation at the expense of neurogenesis during development ^74^. ATF3 is involved in regulating cell fate decisions of radial glia during cortical development ^75^. We see decreased levels of ATF3 in MG following inhibition of NFkB in *Ascl1*-overexpressing mice as well as in MG with deletion of *Ikkb*. Interestingly, ATF3 has been shown to enhance TGFb-signaling in cancer cells^76^, Smad3 has been shown to induce ATF3 upstream of Id1^77^ and TGFb/Smad3 activates ATF3 in hepatic stellate cells during the process of liver fibrosis^78^. Collectively, these findings suggest that ATF3, TGFb2/Smad3 and Ids may coordinate a stress response program down-stream of NFkB and this network interferes with neurogenic programs driven by Ascl1, and promotes glial reactivity and return to quiescence TGFb/Smad3 may be another important signaling hub that coordinates with NFkB to suppress the reprogramming of mammalian MG. We find that inhibition of NFkB or cKO of *Ikkb* results in diminished levels of *Tgfb2*, and inhibition of Smad3 significantly increased in the formation of bipolar-like neurons derived from Ascl1-overexpressing MG, similar to inhibition of NFkB. TGFb/Smad3 signaling has been shown to promote inflammation, scar formation, and fibrosis^79^. In the retina, TGFb signaling through TGFbR2 drives increased expression of GFAP in MG, a symptom of glial reactivity^80^, and TGFb1/2 promotes a gliotic gene networks in the mammalian retina after damage^55^. We find a significant reduction in *Gfap* following inhibition of NFkB, and this may be caused indirectly by diminished TGFb signaling. Additionally, TGFb-Smad3 signaling represses proliferation of MG in the zebrafish retina and promotes a gliotic response following light damage^53^. By comparison, in the chick retina TGFb/Smad2/3 signaling acts in opposition to BMP/Smad/1/5/8 signaling to suppress the formation of MGPCs^56^. Our data indicate that NFkB and TGFb signaling coordinate to promote inflammation/reactive gene networks that suppresses neurogenic reprogramming in the mammalian retina.

Previous studies have shown that Ascl1-overexpressing MG that fail to differentiate into neurons maintain higher expression levels of Ids compared to those cells that acquire neuron-like phenotype^22^. Ids can bind to pro-neural bHLH transcription factors to prevent expression of neuronal genes^58^, while promoting expression of the pro-glial bHLH transcription factor Hes1 to prevent neurogenesis from neural stem cells^51^. Unlike MG in fish and chicks, mammalian MG rapidly undergo a transition to a reactive state after damage to restore quiescence^13^, and expression of Ids may be, in part, responsible for this inability to proceed to neurogenic reprogramming. We find that inhibition of NFkB diminishes expression of *Id1/3*, which may allow for increased neurogenic reprogramming by reducing the inhibition pro-neural gene networks. It has been shown that overexpression of *Id1* can drive degradation of Ascl1^81^, thus MG that maintain Id expression may fail to differentiation into neurons because of Id-mediated degradation of Ascl1. Consistent with these observations, we found that inhibition of NFkB decreased expression of Ids and increased neuronal differentiation with Ascl1 overexpression.

Various inflammatory pathways converge on Id family members. For example, TGFb can induce expression of Id1^82^, and IL-1b and IL6 can induce expression of Id2^83^. We find a significant reduction in expression of both *Il1b* and *Tgfb2* upon inhibition of NFkB signaling, which may be responsible for the reduction of *Id* expression. Collectively our findings suggest that NFkB coordinates with TGFb signaling to promote pro-inflammatory gene networks that converge on pro-glial transcription factors, including NFIs and Ids. Inhibition of NFkB alleviates pro-inflammatory signaling and diminishes expression of pro-glial transcription factors, thereby allowing MG to transition to neurogenic reprogramming rather than restoring quiescence and maintenance of glial phenotype.

## Conclusions

Our findings indicate that NFkB is an important signaling hub that is activated in MG in response to damage. The activation of NFkB in MG relies on signals derived from reactive microglia and regulates the recruitment of peripheral immune cells into damaged retinas. Importantly, our findings indicate that NFkB-signaling, TGFb/Smad-signaling, and Id transcription factors suppress the capacity of MG to de-differentiate and reprogram into neurons with over expression of Ascl1. NFkB-signaling coordinates pro-glial and pro-inflammation gene networks that represses reprogramming, promotes reactivity and drives a return to quiescence.

## Materials and Methods

### Animals

The use of animals in these experiments was in accordance with the guidelines established by the National Institutes of Health and the IACUC committee at the Ohio State University. Mice were kept on a cycle of 12 hours light, 12 hours dark (lights on at 8:00 AM). NFkB-eGFP reporter mice, which have eGFP-expression driven by a chimeric promoter containing three HIV-NFkB cis elements^25^ and *Ikkb^fl/fl^* mice, with insertion of Cre recombinase binding sites (LoxP) into the intronic regions flanking exon 3 of the wild type *Ikkb* gene^32^ were kindly provided Dr. Denis Guttridge’s laboratory at The Ohio State University. We crossed *Ikkb^fl/fl^* mice onto *Rlbp1-CreERT;R26-stop-flox-CAG-tdTomato* mice (provided by Dr. Ed Levine; Vanderbilt University), wherein Cre-mediated recombination occurs in a tamoxifen-dependent manner specifically in MG under the control of the retinaldehyde binding protein 1 (*Rlbp1*) promoter, herein referred to as (*Rlbp1-CreERT:Ikkb^fl/fl^*). The use of Ascl1 over-expressing mice (*Glast-CreER:LNL-tTA:tetO-mAscl1-ires-GFP)* was as previously described^84^.

### Injections

Mice were anesthetized via inhalation of 2.5% isoflurane in oxygen and intraocular injections performed as described previously^85^. For all experiments, the vitreous chamber of right eyes of mice were injected with the experimental compound and the contra-lateral left eyes were injected with a control vehicle. Compounds used in these studies included N-methyl-D-aspartate (NMDA; Sigma; 1.5 ul at concentration of 100mM in PBS), 15-Deoxy-delta12,14-prostaglandin J2 (PGJ2; Alfa Aesar; 5.0 ug/dose in DMSO), TSA (Sigma; 1.0 ug/dose in DMSO), Sis3 (Sigma; 2.5 ug/dose in DMSO), AGX51 (Fischer; 10.0 ug/dose in DMSO). Intraperitoneal injections of tamoxifen (Sigma; 1.5 mg/100 μl corn oil per dose) were performed for 4 consecutive days to induce ER-Cre activity.

### Single Cell RNA-sequencing of retinas

Retinas were obtained from adult (age ≥ postnatal day 60) mouse retinas at various times after NMDA treatment. Retinas were dissociated in papain/DNase (Worthington Biochemical) for 15 minutes at 37°C then triturated to form single cell suspensions. Ovomucoid (Worthington Biochemical) was added to halt the proteolysis and samples were centrifuged at 300xG for 10 minutes at 4°C and resuspended in a 1:1:10 solution of DNase:ovomucoid:Neurobasal media (Worthington:Worthington:Gibco). Cell suspensions were incubated with primary antibodies to CD11b and CD45 (**table 1**) at a dilution of 1:1000 in DNase:ovomucoid:Neurobasal (1:1:10) for 15 minutes. Cells were sedimented and rinsed in DNase:ovomucoid:Neurobasal (1:1:10) twice for 10 minutes. The cell suspension was passed through a 40 µm nylon filter. FACs was performed on a BD FACS ARIAIII Cell Sorter for GFP^+^ MG or TdTomato^+^ MG and CD45^+^/Cd11b^+^ immune cells (Flow Cytometry Shared Resource core at OSU).

Following FACS, cell suspensions were assessed for viability (Countess II; Invitrogen) and cell-density diluted to 700 cell/µl. Each single cell cDNA library was prepared for a target of 10,000 cells per sample. Cells and 10X Genomics Chromium Single Cell 3’ V3 reagents were loaded onto chips to capture cells with gel beads in emulsion (GEMs) using 10X Chromium Controller. Library preparation was according to manufacturer’s protocols. Sequencing was conducted using an S4 flowcell and NovaSeq 6000 (Novogene) with the following parameters: Read 1 i7 Index i5 Index Read 2 28 cycles 8 cycles 0 cycles 91 cycles. Fasta sequence files were de-multiplexed, aligned and annotated to the mm10 genome using 10X Genomics Cell Ranger software.

Using Seurat toolkits^86,87^,Uniform Manifold Approximation and Projection (UMAP) for dimensional reduction plots were generated from aggregates of multiple scRNA-seq libraries. Seurat was used to construct gene lists for differentially expressed genes (DEGs), violin/scatter plots and dot plots. Significance of difference in violin/scatter plots was determined using a Wilcoxon Rank Sum test with Bonferroni correction. Monocle was used to construct unguided pseudotime trajectories and scatter plotters for MG and MGPCs across pseudotime^88–90^. SingleCellSignalR was used to assess potential ligand-receptor interactions between cells within scRNA-seq datasets^45^. Lists of DEGs were uploaded to ShinyGo v0.66 (http://bioinformatics.sdstate.edu/go/) and AmiGO 2 (http://amigo.geneontology.org/amigo) to perform Gene Ontology (GO) enrichment analyses.

Genes that were used to identify different types of retinal cells included the following: (1) Müller glia: *Glul, Nes, Vim, Scl1a3, Rlbp1*, (2) microglia: *C1qa, C1qb, Csf1r, Apoe*, *Aif1* (3) ganglion cells: *Thy1, Pou4f2, Rbpms2, Nefl, Nefm*, (4) amacrine cells: *Gad67, Calb2, Tfap2a*, (5) horizontal cells: *Prox1, Calb2*, (6) bipolar cells: *Vsx1, Otx2, Grik1, Gabra1*, and (7) cone photoreceptors: *Gnat2, Opn1lw*, and (8) rod photoreceptors: *Rho, Nr2e3, Arr1*.

### Fixation, sectioning and immunocytochemistry

Tissues were fixed in 4% PFA, sectioned, and immunolabeled as described previously^91^. Working dilutions and sources of antibodies used in this study are listed in **table 1**. None of the observed labeling was due to non-specific labeling of secondary antibodies or autofluorescence because sections labeled with secondary antibodies alone were devoid of fluorescence. Secondary antibodies included donkey-anti-goat-Alexa488/568, goat-anti-rabbit-Alexa488/568/647, goat-anti-mouse-Alexa488/568/647, goat anti-rat-Alexa488 (Life Technologies) diluted to 1:1000 in PBS plus 0.01% Triton X-100, and incubated for 1 hour.

### Labeling for EdU

Regular drinking water was removed 24hr prior to NMDA injections and replaced with water containing 5-ethynyl-2′-deoxyuridine (EdU; Sigma; 50 mg/100 mL dH2O) and EdU water was replaced every third day. Mice were maintained on EdU water until 4^th^ day after TSA treatment.

Immunolabeled tissue sections were fixed in 4% formaldehyde in PBS for 5 minutes at room temperature, washed for 10 minutes with PBS, permeabilized with 0.01% Triton X-100 in PBS for 1 minute at room temperature, and washed in PBS for 10 minutes. Sections were incubated for 30 minutes at room temperature in 2M Tris, 50 mM CuSO_4_, Alexa Fluor 568 or 647 Azide (Thermo Fisher Scientific), and 0.5M ascorbic acid in dH_2_O. Sections were washed with PBS and further processed for immunofluorescence as required.

### Terminal deoxynucleotidyl transferase dUTP nick end labeling (TUNEL)

To identify dying cells that contained fragmented DNA the TUNEL assay was used. We used an *In Situ* Cell Death Kit (TMR red; Roche Applied Science), as per the manufacturer’s instructions.

### Photography, immunofluorescence measurements, and statistics

Wide-field photomicroscopy was performed using a Leica DM5000B microscope equipped with epifluorescence and Leica DC500 digital camera or Zeiss AxioImager M2 equipped with epifluorescence and Zeiss AxioCam MRc. Confocal images were obtained using a Leica SP8 imaging system at Department of Neuroscience Imaging Facility at The Ohio State University. Images were optimized for color, brightness and contrast, multiple channels overlaid and figures constructed by using Adobe Photoshop. Cell counts were performed on representative images. To avoid the possibility of region-specific differences within the retina, cell counts were consistently made from the same region of retina for each data set.

Similar to previous reports^65,92^, immunofluorescence was quantified by using Image J (NIH). Identical illumination, microscope, and camera settings were used to obtain images for quantification. Retinal areas were sampled from 5.4 MP digital images. Measurements of fluorescence of ATF3 within the nuclei of MG were made by selecting the total area of pixel values ≥70 for Sox2 immunofluorescence (in the red channel) and copying the fluorescence of ATF3 within this area in the green channel. This copied channel data was pasted into a separate file for quantification or onto 70% grayscale background for figures. Measurements were made for regions containing pixels with intensity values of 68 or greater (0 = black and 255 = saturated). The total area was calculated for regions with pixel intensities >68. The average pixel intensity was calculated for all pixels within threshold regions. The density sum was calculated as the total of pixel values for all pixels within threshold regions. These calculations were determined for retinal regions sampled from ≥6 different retinas for each experimental condition. The mean area, intensity and density sum was calculated for the pixels within threshold regions from ≥6 retinas for each experimental condition.

Where significance of difference was determined between two treatment groups accounting for inter-individual variability (means of treated-control values) we performed a two-tailed, paired t-test. Where significance of difference was determined between two treatment groups, we performed a two-tailed, unpaired t-test. Where evaluating significance in difference between multiple groups we performed ANOVA followed by Tukey’s test. GraphPad Prism 6 was used for statistical analyses and generation of histograms and bar graphs.

## Data availability

scRNA-seq data: GSE135406

Some scRNA-seq libraries can be queried at: https://proteinpaint.stjude.org/F/2019.retina.scRNA.html and gene-cell matricies can be downloaded from GitHub at https://github.com/fischerlab3140/scRNAseq_libraries

## Acknowledgements

This work was supported by RO1 EY022030-08 (AJF), UO1 EY027267-04 (SB, AJF), and T32 NS105864 (IP). We thank Dr. Ed Levine for providing the *Rlbp1-*CreERT mice and Dr. Dennis Guttridge for providing the NFkB-eGFP reporter mice and *Ikkb^fl/fl^* mice. We thank Heithem El-Hodiri and Olivia Taylor for comments and discussions that contributed to the final form of the paper.

## Author Contributions

IP designed and executed experiments, gathered and analyzed data, constructed figures and wrote the manuscript. TH and SB generated some of the scRNA-seq databases. LT and TR consulted on experimental design and provided the Ascl1 over-expressing mice. AJF designed experiments, analyzed data, constructed figures and wrote the manuscript.

## Supplemental Figure Legends

**Supplemental Fig. 1:**
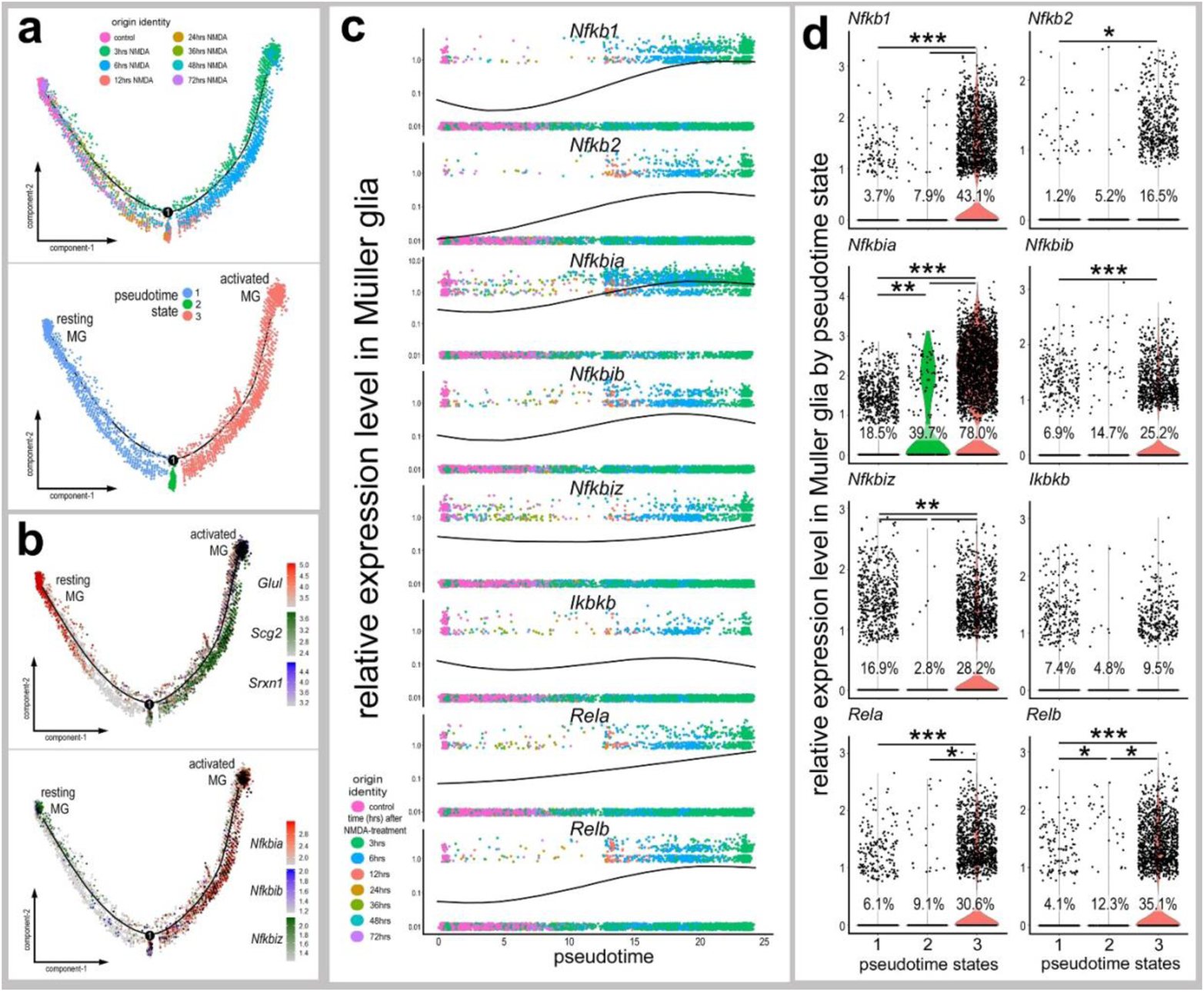
Expression of NFKB factors across pseudotime in damaged Müller glia. Pseudotime trajectory plots of MG from control and 3, 6, 12, 24, 36, 48 and 72h after NMDA damage separate into 3 distinct clusters **(a)**. Resting glia are identified by expression of *Glul*, activated glia are identified by expression of *Scg2* and *Srxn1*. The pseudotime trajectory plots **(b)** demonstrate *Nfkbia, Nfkbib, Nfkbiz* enriched in activated glia. Pseudotime reduction plots **(c)** and violin plots **(d)** illustrate change in expression levels of *Nfkb1, Nfkb2, Nfkbia, Nfkbib, Nfkbiz, Ikbkb, Rela*, and *Relb*. Significance of difference (**P<0.01, ***P<0.001) was determined by using a Wilcoxon rank sum with Bonferroni correction.

**Supplemental Fig. 2:**
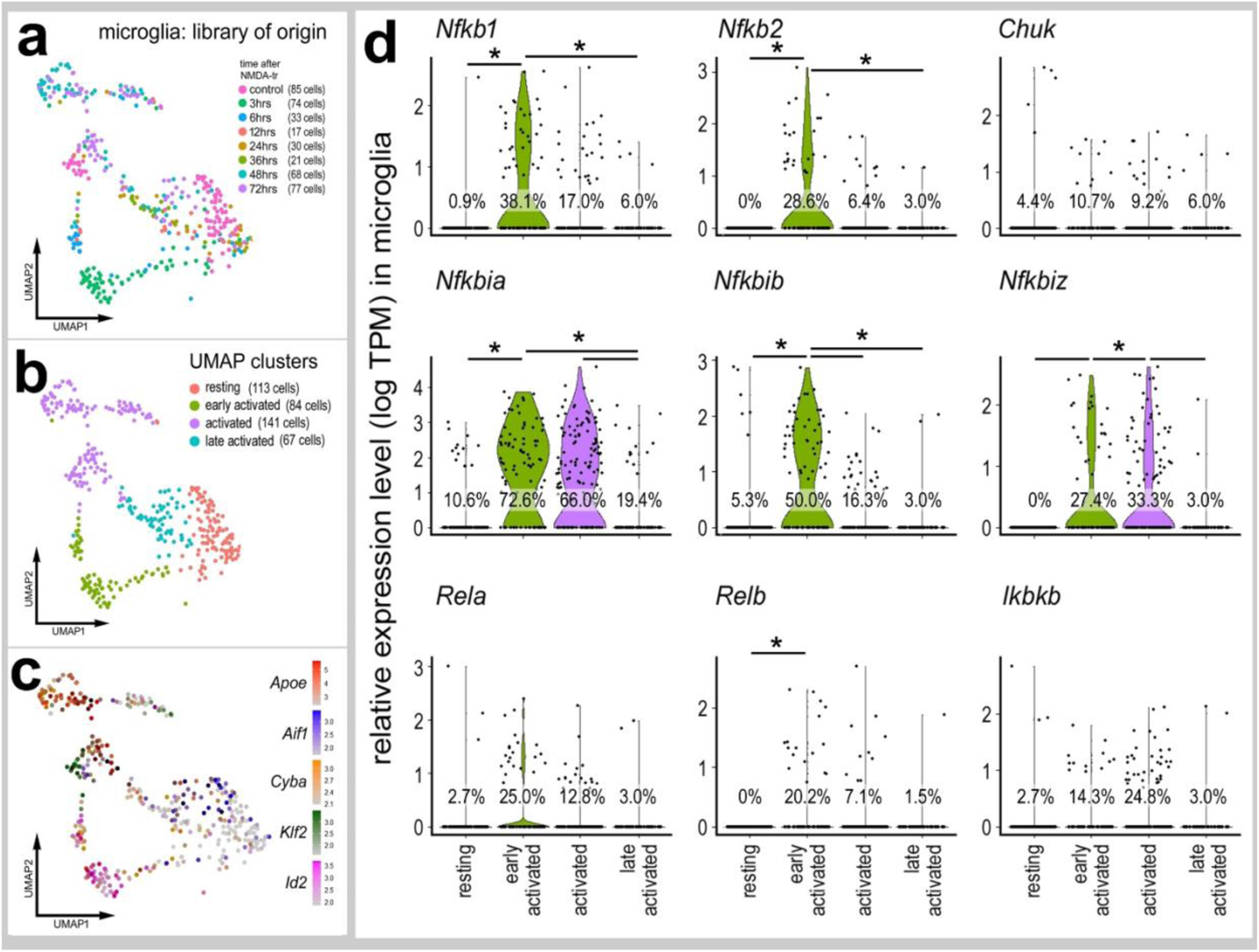
NFkB factors expressed in microglia. UMAP plots of isolated and re-embedded microglia from WT control and NMDA 3, 6, 12, 24, 36, 48 and 72h scRNA-Seq libraries show 4 distinct clusters **(a,b**). Clusters were identified based on expression of *Apoe, Aif1, Cyba, Klf2*, and *Ids* **(c**). Violin plots illustrate levels of expression and percentage of expressing cells for *Nfkb1, Nfkb2, Chuk, Nfkbia, Nfkbib, Nfkbiz, Rela, Relb, Ikbkb* in different UMAP clusters of microglia **(d**). Significance of difference (**p<0.01, ***p<0.001) was determined by using a Wilcoxon rank sum with Bonferroni correction.

**Supplemental Fig. 3:**
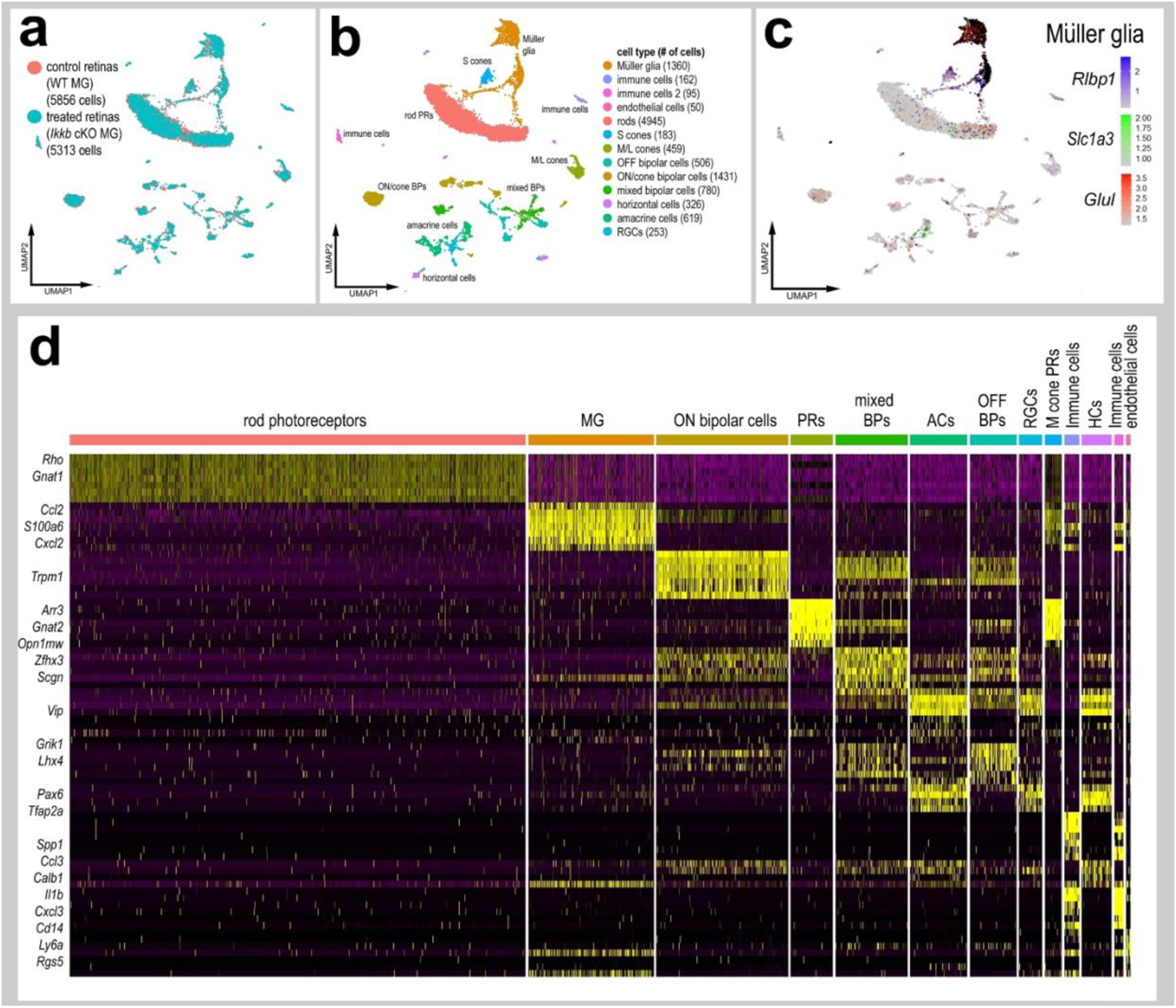
scRNA-Seq for whole retina *Ikkb*-cKO. UMAP plots for whole retina scRNA-seq libraries of *Rlbp1-CreERT* (control) and *Rlbp1-CreERT:Ikkb^fl/fl^* (treated) mice at 8h after NMDA damage **(a)**. Distinct cell clusters of UMAP ordered cells **(b)** were characterized based on expression of markers such as *Rlbp1, Slc1a3* and *Glul* **(c)** and DEGs identified from heatmap plots **(d)**.

**Supplemental Fig. 4:**
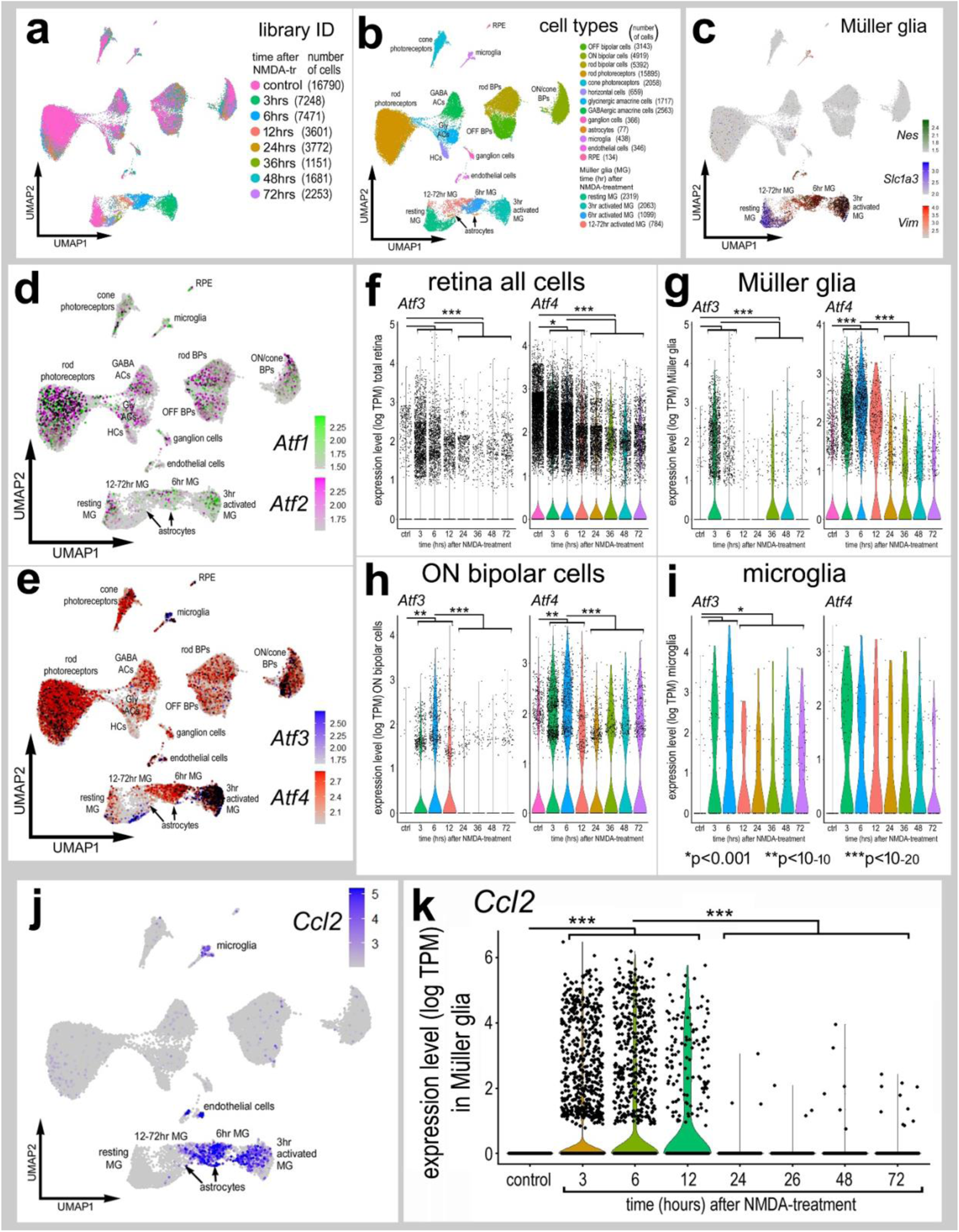
Expression patterns of *Atf’*s and *Ccl2* in WT scRNA-Seq libraries from control and damaged retinas. Aggregated scRNA-seq libraries were prepared from WT control retinas and retinas 3, 6,12, 24, 36, 48 and 72h after NMDA damage **(a)**. Clusters were identified based on expression of cell-distinguishing markers described in Methods **(b)**. Resting MG and reactive MG identified by expression of *Slc1a3* or *Nes and Vim*, respectively **(c)**. UMAP plots illustrate patterns of expression of *Atf1* and *Atf2* **(d),** *Atf3* and *Atf4* **(e)**, or *Ccl2* **(j)** across retinal cell types. Violin plots illustrate expression levels of *Atf3* and *Atf4* in MG, bipolar cells, and microglia **(f-i**), or *Ccl2* in MG **(k)**. Significance of difference (**p<0.001, **p<1×10-10, ***p<1×10-20) was determined by using a Wilcoxon rank sum with Bonferroni correction.

**Supplemental Fig. 5:**
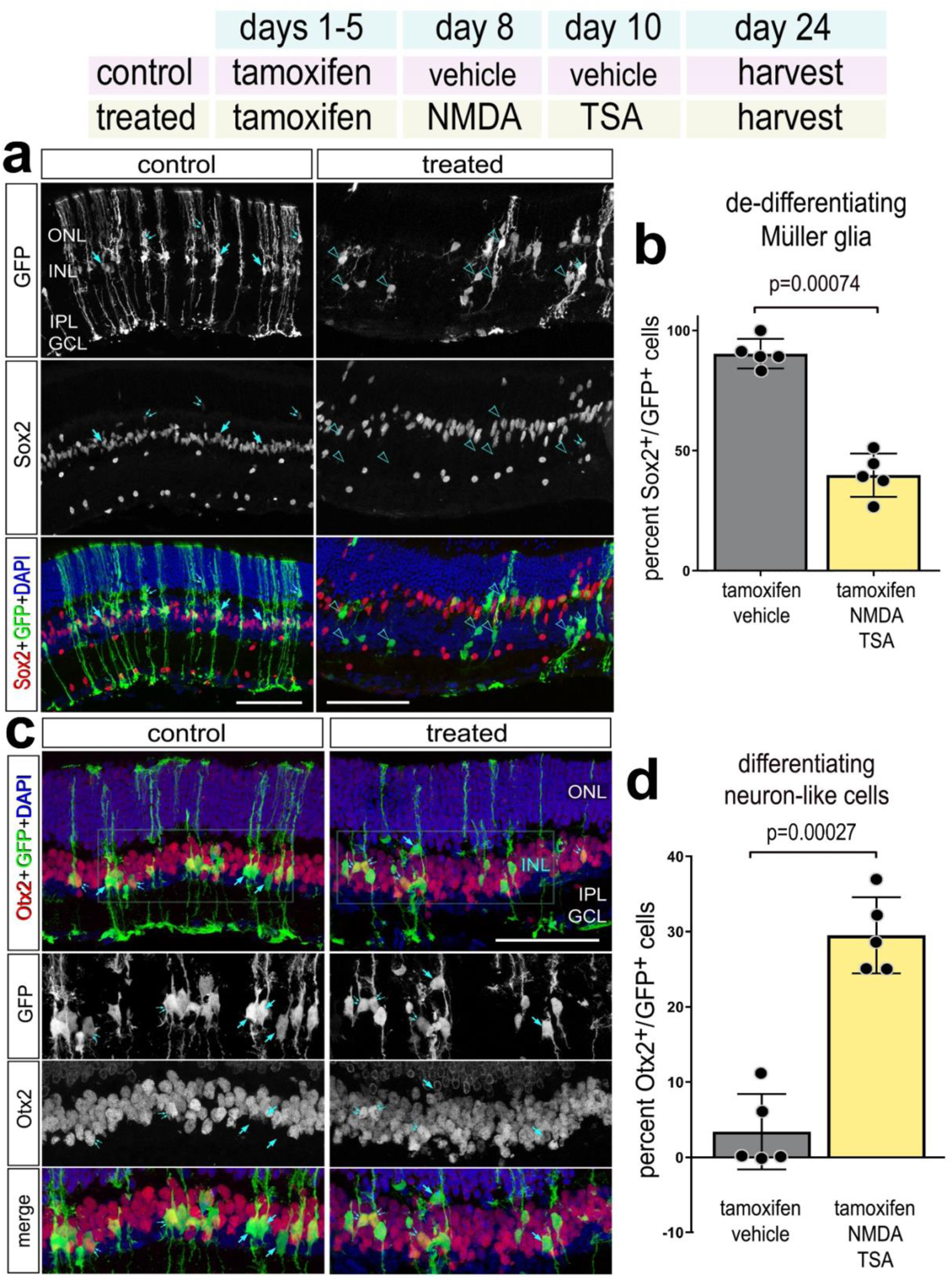
Validation of ANT regeneration paradigm. Experimental paradigm outlined at top. Retinal sections were labeled for GFP (green), DAPI (blue), Sox2 (red; **a**), or Otx2 (red; **c**). Arrows indicate double-labeled cells. Histograms illustrate the mean percentage (±SD and individual data points) of GFP^+^/Sox2^+^ **(b)** or GFP^+^/Otx2^+^ **(d)** cells. Significance of difference (p-values shown) was determined by using a paired t-test.

**Supplemental Fig. 6:**
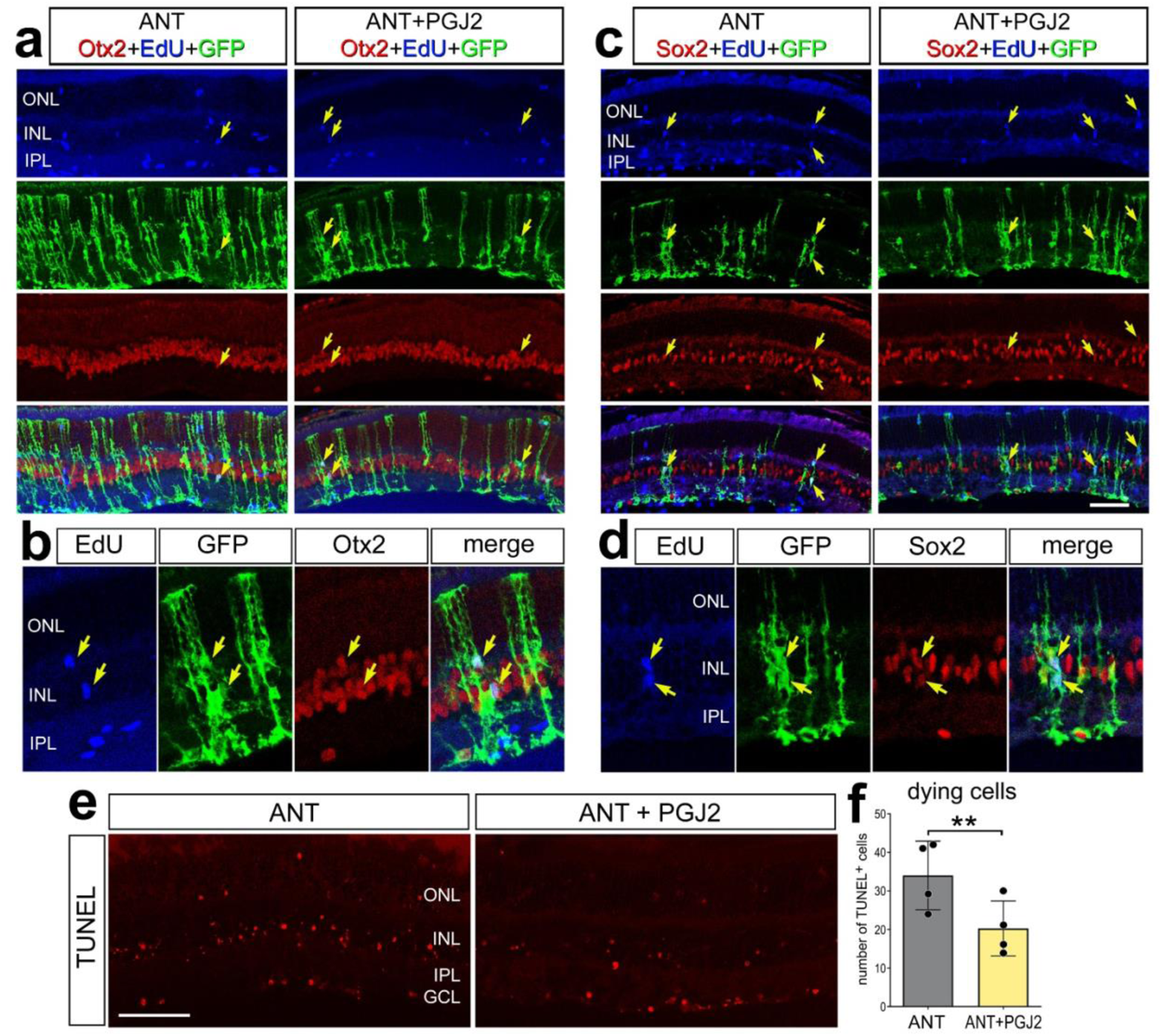
Inhibition of NFkB signaling promotes proliferation and neuroprotection following ANT treatment. Experimental paradigm outlined at top. Tamoxifen was administered IP 1x daily for 4 consecutive days in *Glast*-*CreER:LNL-tTA:tetO-mAscl1-ires-GFP* mice, NMDA was injected intravitreally in left (control) eyes or NMDA+PGJ2 in right (treated) eyes on D8, vehicle ± PGJ2 on D9, TSA ± PGJ2 on D10, and retinas were harvested 4 days after TSA. EdU was added to the drinking water and administered starting on D7 and sustained until harvesting retinas on D14. Retinal sections were labeled for GFP (green), Otx2 (red;**a-b**), Sox2 (red;**c-d**), and EdU (blue). Arrows in **a** and **b** indicate GFP^+^/EdU^+^/Otx2^+^ cells, and arrows in **c** and **d** indicate GFP^+^/EdU^+^/Sox2^+^ cells. TUNEL assays were performed on ANT or ANT+PGJ2 treated retinas at 24h after NMDA **(e)**. Histograms illustrate the mean (±SD and individual data points) number of TUNEL^+^ cells **(f)**. Significance of difference (**p<0.01) was determined by using a paired t-test.

**Supplemental Fig. 7:**
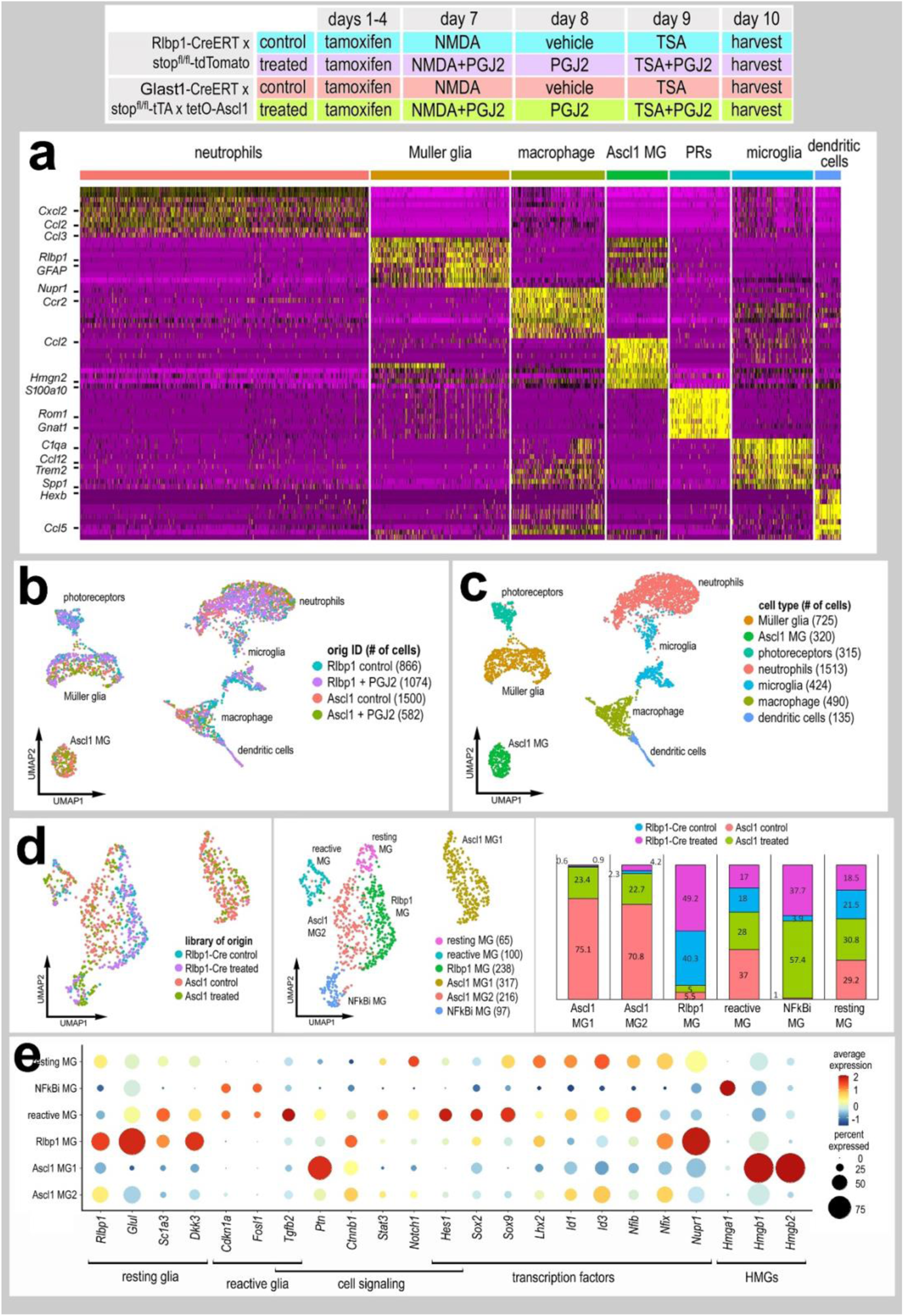
scRNA-Seq libraries from Ascl1 overexpressing mice. scRNA seq libraries were prepared from *Rlbp1-CreERT* and *Glast*-*CreER:LNL-tTA:tetO-mAscl1-ires-GFP* mice. Tamoxifen was injected IP 1x daily for 4 consecutive days. NMDA was injected intravitreally into left (control) eyes and NMDA+PGJ2 in right (treated eyes) on D8, vehicle ± PGJ2 on D9, TSA ± PGJ2 on D10, and retinas were harvested for scRNA-seq library prep 24h after TSA treatment. Samples were enriched via FACS for either GFP^+^ cells and CD45^+^/Cd11b^+^ cells (from Ascl1 mice) or TdTomato^+^ cells and CD45^+^/Cd11b^+^ cells (from *Rlbp1-CreERT* mice). Clusters of different types of retinal cells were identified based on expression patterns of different cell-distinguishing markers **(a). (b-c)** UMAP plots of aggregated scRNA-seq libraries for MG and immune cells. **(d)** UMAP plots of isolated and re-embedded MG from *Rlbp1-CreERT* control (NMDA) and treated (NMDA+PGJ2) and Ascl1 control (NMDA) and treated (NMDA+PGJ2). Stacked bar graphs illustrate UMAP cluster occupancy by library origin identity **(d)**. The dot plot in **(e)** illustrates changes in expression of resting and reactivity genes, cell signaling factors, and transcription factors. Dot size represents percent of cells expressing and dot color representing expression levels.

